# The X-linked splicing regulator MBNL3 has been co-opted to restrict placental growth in eutherians

**DOI:** 10.1101/2021.09.08.459166

**Authors:** Thomas Spruce, Mireya Plass, André Gohr, Debashish Ray, María Martínez de Lagrán, Gregor Rot, Ana Nóvoa, Demian Burguera, Jon Permanyer, Marta Miret, Hong Zheng, Maurice S. Swanson, Quaid Morris, Moises Mallo, Mara Dierssen, Timothy R. Hughes, Barbara Pernaute, Manuel Irimia

**Author notes:** Corresponding authors. **Thomas Spruce**, Centre for Genomic Regulation, Dr. Aiguader, 88, 08003 Barcelona, Spain, **Barbara Pernaute**, Centre for Genomic Regulation, Dr. Aiguader, 88, 08003 Barcelona, Spain, **Manuel Irimia**, Centre for Genomic Regulation, Dr. Aiguader, 88, 08003 Barcelona, Spain.

## Abstract

The eutherian placenta is a major site for parental genetic conflict. Here, we identify the X-linked *Mbnl3* gene as a novel player in this dispute. *Mbnl3* belongs to an RNA binding protein family whose members regulate alternative splicing and other aspects of RNA metabolism in association with cellular differentiation. We find that, in eutherians, *Mbnl3* has become specifically expressed in placenta and has undergone accelerated sequence evolution leading to changes in its RNA binding specificities. Although its molecular roles are partly redundant with those of *Mbnl2, Mbnl3* has also acquired novel biological functions. In particular, whereas *Mbnl2*;*Mbnl3* double knockout mice display severe placental maturation defects leading to strong histological and functional abnormalities, *Mbnl3* knockout alone results in increased placental growth and favors placental and fetal resource allocation during limiting conditions.

## Introduction

Placentas have evolved independently many times in vertebrates ^1^ and provide a direct link between mother and fetus through which nutrient exchange can occur. In eutherians, the placenta is mainly formed from the trophoblast, a mammalian specific innovation derived from the embryo in the first lineage decision of development. Placental evolution results in the loss of full maternal control of fetal resource allocation via yolk formation. In species where females breed with multiple males over their lifetimes this may result in the placenta becoming a site of parental conflict over resource allocation ^2^. Such conflicts are proposed to occur because, in those species, siblings are more likely to be related via their maternal than their paternal genome. This means that the inclusive fitness of paternally inherited alleles is optimal when there is a high level of resource allocation to the individual offspring containing them. On the other hand the inclusive fitness of maternally inherited alleles is optimized when there is a greater degree of maternal resource sharing between siblings and thus lower levels of maternal investment in individual offspring ^3^. Genetic conflicts over resource allocation have likely contributed to the accumulation of genes with placentally biased expression on the X-chromosome ^4^ as sex chromosomes are expected to accumulate sexually antagonistic genes due to their asymmetric inheritance pattern and distribution ^5-7^. This effect is likely enhanced in species such as mouse, where trophoblastic gene expression from the X-chromosome is strongly paternally imprinted and occurs almost exclusively from maternal alleles ^8^, which is likely to drive the accumulation of maternally favorable variants reducing resource allocation ^9-11^. Indeed, in mouse, placental hyperplasia is seen when a number of placentally expressed X-linked genes are knocked out ^12-14^.

To date, understanding the regulation of protein production in the eutherian placenta has mainly been focused at the transcriptional level ^15,16^. In contrast, with the exception of miRNA-mediated modulation of gene expression ^17^, little is known about the role of post-transcriptional regulation of processes such as alternative splicing, alternative polyadenylation and mRNA translation. In particular, alternative splicing, in which an exon or intron is either excluded from or included in a transcript, has the potential to greatly expand both transcriptomic and proteomic diversity ^18-20^. RNA binding proteins mediate much of this post-transcriptional regulation, with the same proteins often regulating multiple aspects of RNA metabolism ^21,22^. Targeting of RNA by these regulators is usually mediated by RNA binding domains ^23^, which are typically discrete and ordered. However, it has recently become apparent that intrinsically disordered regions can also mediate binding ^24^. RNA binding proteins often contain multiple RNA binding domains, increasing both their binding specificity and affinity ^22^.

To find post-transcriptional regulators with relevance for placenta development and evolution, we first searched for splicing regulators with enriched expression in trophoblastic vs. non– trophoblastic tissues. We identified two Muscleblind-like (MBNL) genes, *Mbnl2* and *Mbnl3*, that through a combination of comparative genomic approaches and *in vitro* binding assays we found to have undergone strikingly different evolutionary paths during the emergence of the eutherian lineage. Using mouse knockout approaches coupled with high throughput sequencing, we investigated the functional role of these factors in placenta and identified a unique role in restricting placental growth for the X-linked gene *Mbnl3*.

## Results

### Mbnl3 *has the strongest placenta enrichment among splicing factors*

In order to identify splicing regulators with a potential placenta-specific function, we began by screening a combination of publicly available and in-house transcriptomic datasets (Supplementary Table 1) for genes that are differentially expressed between trophoblastic and embryonic tissues in mouse. The following expression comparisons were made: (i) placenta vs. non-placental tissues, (ii) trophoblast vs. inner cell mass at the blastocyst stage, and (iii) trophoblast stem cells vs. embryonic and extraembryonic endoderm stem cells (Fig. 1a-c and Supplementary Table 2; see Methods). These analyses revealed the X-linked gene *Mbnl3* to be substantially more enriched than any of the other regulators examined in both placenta (analysis (i)) and E3.5 trophoblast (analysis (ii)) (Fig. 1a,b). Six other genes were also identified as being enriched (log2FC > 1) in trophoblastic tissues in at least one of these comparisons: *Mbnl2* (an autosomal *Mbnl3* paralog), *Esrp1, Esrp2, Rbm20, Rbm38* and *Rbms1*. Expression enrichment in the trophoblast for the top candidates was confirmed by *in situ* hybridization analysis (Fig. 1d and Extended Data Fig. 1a). This showed expression of *Mbnl3* and *Esrp2* to be restricted to differentiated trophectodermal tissue, *Rbm20* to be specifically expressed in the trophoblast stem cell compartment, and *Mbnl2, Rbm38* and *Rbms1* to be expressed in both the trophoblast stem cell compartment and more differentiated trophectodermal tissues. Importantly, of these splicing factors, only *Mbnl3* was highly placenta specific, with no or low expression in other tissues (Fig. 1e and Extended Data Fig. 1e). Moreover, analysis of RNA-seq data from human and cow showed placental expression of these factors to be highly conserved, with *Mbnl3* also showing the strongest placenta-specific enrichment alongside very low levels of expression in non-placental tissues (Fig. 1f, Extended Data Fig. 1c-e and Supplementary Tables 1,3,4). In contrast, *Mbnl3* was strongly expressed across a range of adult tissues in non-eutherians (Extended Data Fig. 1e and Supplementary Table 1), including the marsupial opossum, for which *Mbnl3* showed no placenta-enriched expression (Fig. 1f and Supplementary Table 5).

**Fig. 1.**
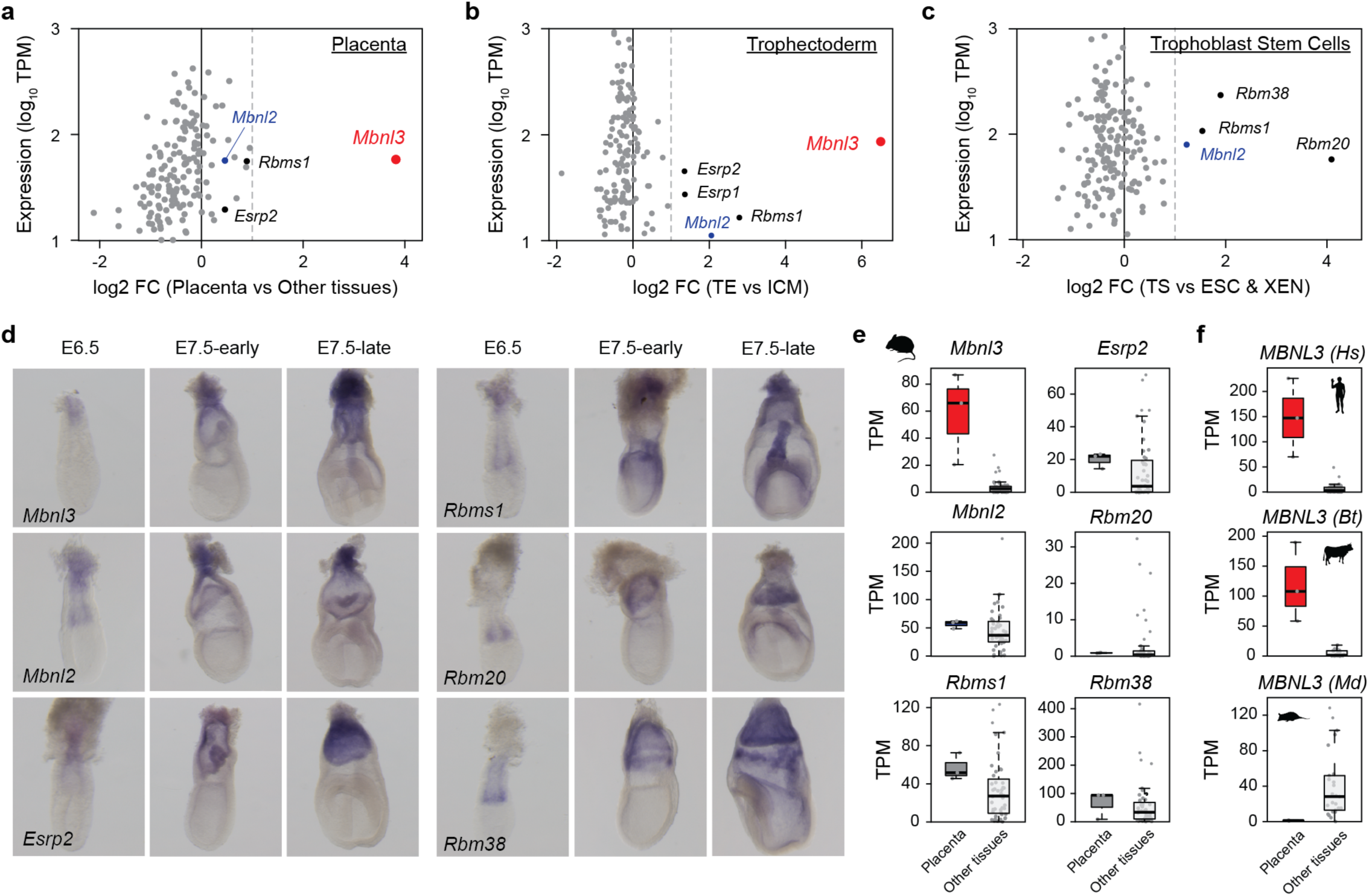
Analysis of splicing factor expression enrichment in trophoblastic vs non-trophoblastic tissues. **a-c**, Scatter plots showing trophoblastic expression level and enrichment for 197 splicing regulators in three separate comparisons of trophoblastic vs non-trophoblastic tissues. **d**, Whole mount *in situ* hybridisation analysis of the expression of splicing regulators found to be enriched in trophoblastic tissues (a-c) in mouse embryos at the indicated stages. **e**,**f**, Box plots showing expression levels of the indicated splicing factors in placenta and non-placental tissues in mouse (e) and in human (Hs), cow (Bt), and opossum (Md)(f).

### *Placental expression of* Mbnl3 *from a novel eutherian-specific promoter*

Comparison of the *Mbnl3* gene locus across a number of vertebrate species revealed it harbors a novel eutherian specific transcriptional start site (TSS), adjacent to a eutherian-specific hAT family DNA-transposon *MER53*, in addition to the canonical TSSs, conserved across vertebrates, and more lineage-restricted TSSs (Fig. 2a and Extended Data Fig. 2a-c). The use of the eutherian-specific TSS, or of rodent or primate specific TSSs found within < 5 kbp of it, results in a transcript that codes for a N-terminally truncated protein (Fig. 2b). A similarly truncated MBNL3 protein isoform can also be generated from alternatively spliced transcripts that initiate from the ancestral TSSs but skip exon 2, which encodes the canonical start codon (Fig. 2a). Consistent with previous studies, we found that both full-length and truncated MBNL3 protein isoforms were present in mouse placenta by Western blot analysis (Extended Data Fig. 2d). To investigate the use of the different TSSs and N-terminal isoforms, we quantified RNA-seq reads spanning competing exon-exon junctions (A-D in Fig. 2a) across a range of tissues from mouse, human and cow (Fig. 2c, Extended Data Fig. 2e,f and Supplementary Table 6). This analysis revealed the eutherian-specific TSS, alongside the nearby lineage-specific ones, to be dominant in blastocyst-stage trophoblast tissue in all three species. Furthermore, in placenta, the eutherian-specific TSS was found to account for around half of the transcripts in mouse and almost all of the transcripts in human and cow. This strong trophoblastic usage of the eutherian-specific TSS prompted us to test the ability of the region surrounding it to act as enhancers/promoters by transgenic reporter assay. This revealed these regions to contain regulatory elements capable of driving trophoblast-specific expression in mouse embryos (Fig. 2d).

**Fig. 2.**
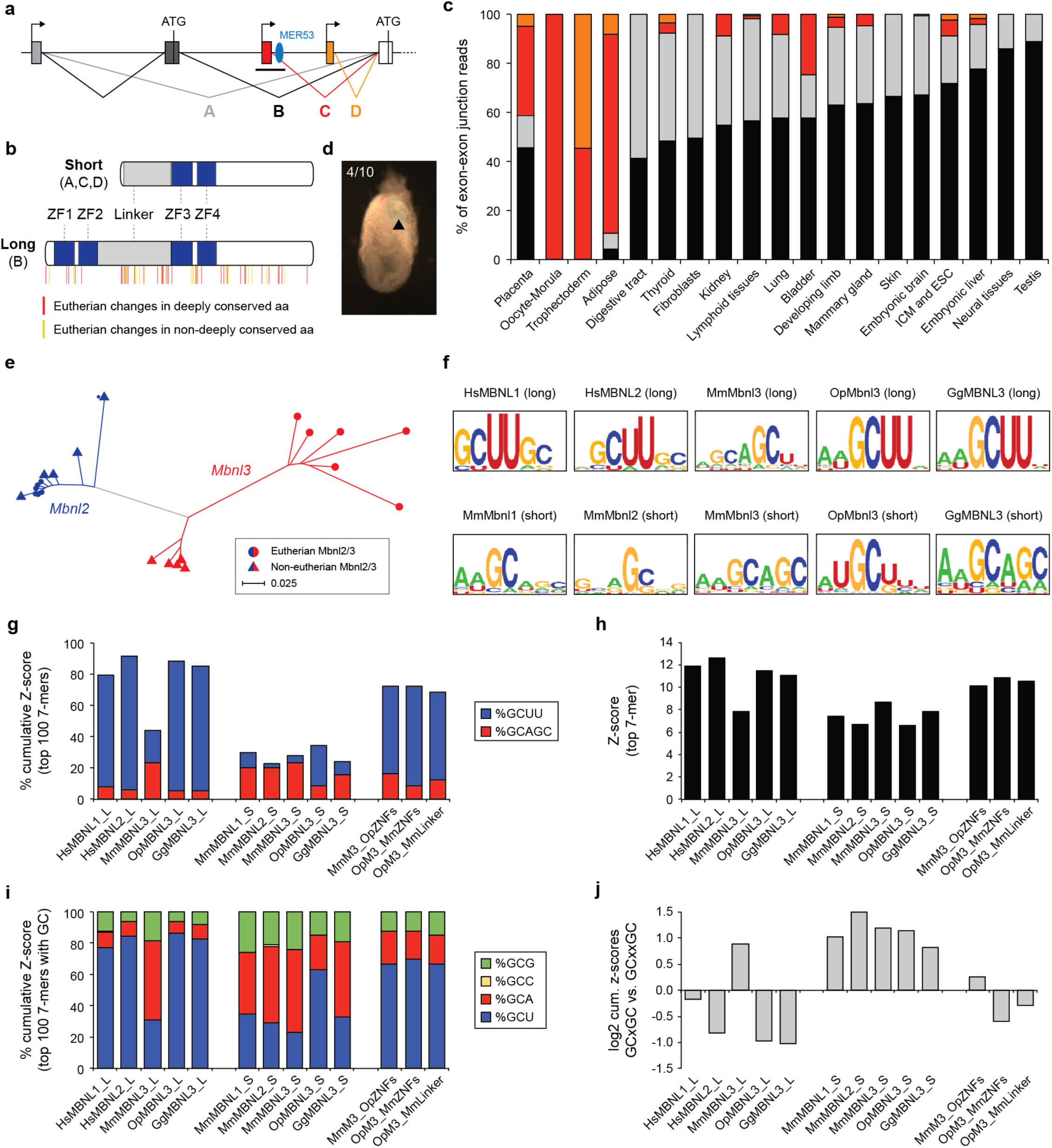
Molecular evolution of *Mbnl3* in eutherians. **a**, Schematic showing alternative splicing events and transcriptional (arrow) and translational (ATG) start sites leading to long and short MBNL3 isoform production. Splice junctions corresponding to transcripts that code for the long (B) and short (A,C,D) isoforms are indicated. Ancestral (black/grey), eutherian-specific (red) and mouse-specific (orange) transcriptional start sites and the location of the DNA-transposon MER53 (blue) are also shown. **b**, Diagram showing long and short Mbnl3 isoforms and the location of eutherian-specific amino acid (aa) changes. Zinc finger domains are shown in blue. Sequence alignment is shown in Extended Data Fig. 3. **c**, Quantification of the relative usage of the splice junctions highlighted in (a) across a range of mouse tissues. **d**, Transgenic analysis of a putative Mbnl3 promoter region. A ∼8kb region surrounding the eutherian specific transcriptional start site indicated by the black bar in (a) produced lacZ staining in the trophoblast of 4/10 injected embryos at E7.5. **e**, Phylogenetic tree showing the degree of MBNL3 (red) and MBNL2 (blue) protein sequence divergence in eutherian (circles) and non-eutherian (triangles) species. The tree was produced by the neighbour-joining method using Kimura corrected distances. **f**, RNAcompete-derived sequence logos for the indicated human (Hs), mouse (Mm), opossum (Op) and chicken (Gg) MBNL proteins. **g**, Analysis of the Z-score contribution of sequences containing GCUU or GCAGC motifs to the total cumulative Z-score of the top 100 RNAcompete derived 7-mers for the indicated MBNL proteins. **h**, Bar plot showing the Z-score of the top RNA-compete derived 7-mer for each of the indicated MBNL proteins. **i**, Analysis of the Z-score contribution of sequences containing GCG, GCC, GCA or GCU motifs to the total cumulative Z-score of the top 100 RNA compete derived 7-mers containing a single GC dinucleotide for the indicated MBNL proteins. **j**, Ratio of the z-score contribution of 7-mer’s containing a pair of GC dinucleotides with a 1bp spacer (GCxGC) vs a 2bp spacer (GCxxGC).

### MBNL3 has undergone accelerated sequence evolution in eutherians

MBNL proteins contain two pairs of zinc fingers (ZF1-2 and ZF3-4), which mediate RNA binding, separated by a linker region, which is required for splicing regulation and that may play a role in RNA binding (Fig. 2b; ^25-29^). The first pair of zinc fingers is missing from the truncated eutherian MBNL3 isoform (Fig. 2b), and equivalent truncated protein isoforms have not been observed for MBNL1 or MBNL2 ^30-32^. Using phylogenetic analysis, we found that *Mbnl3* has undergone accelerated sequence evolution in eutherians compared to non-eutherian *Mbnl3* and all *Mbnl2* genes (Fig. 2e). In particular, eutherian MBNL3 proteins show specific changes in conserved amino acids in ZF1, the spacers separating ZFs 1 and 2 and ZFs 3 and 4, as well as the linker region (Fig. 2b and Extended Data Fig. 3a). In contrast, the protein sequence of MBNL2 is highly conserved across vertebrates. Close examination of *Mbnl3* gene structure identified an additional feature to be associated with eutherians: loss of a 54 bp alternative exon ^33^, which triggers nuclear localization upon inclusion ^28^ and that is found in all *Mbnl1* and *Mbnl2* paralogs and in most non-eutherian *Mbnl3* genes (Extended Data Fig. 3b). Altogether, these eutherian specific changes to *Mbnl3*, alongside the loss of its expression from non-placental tissues and its location in the X-chromosome, suggest it may have undergone a process of evolutionary specialization in this clade.

### Eutherian specific changes in MBNL3 binding preferences

To investigate how these eutherian-specific novelties may have affected MBNL3 functionality, we first examined the RNA binding preference of full-length (long) eutherian (mouse) and non-eutherian (chicken and opossum) MBNL3 proteins using RNAcompete ^34-36^. We found that full-length opossum and chicken MBNL3 show a strong GCUU binding preference, similar to the binding preferences that have previously been described for mammalian MBNL1 and MBNL2 and non-vertebrate MBNL proteins (Fig. 2f,g)^33,34,37-39^. In contrast, we found full-length mouse MBNL3 to have overall weaker binding preferences, as measured by the top Z-score values (Fig. 2h), and to preferentially bind a GCA/U motif similar to that identified by previous CLIP-seq based studies (Fig. 2f,g,i)^33,40,41^. Furthermore, we observed that amongst 7-mer motifs containing a pair of GC di-nucleotides, opossum and chicken MBNL3 show a strong preference for a pair of uracil’s between the GCs while mouse MBNL3 appears to favor a single adenosine residue in that position (GCUUGC vs GCAGC; Fig. 2g,i,j). Altogether these results demonstrate that the binding specificity of MBNL3 has specifically diverged in eutherians.

Given the high prevalence of the short MBNL3 protein isoform in trophoblastic tissues, we also assessed its binding preferences alongside those of artificially created short isoforms for chicken and opossum MBNL3 and for mouse MBNL1 and MBNL2. The short isoform of mouse MBNL3 was found to have similar GCU/A binding preferences to the long isoform suggesting both may bind the same target sites (Fig. 2f-j). Intriguingly, similar binding preferences were also seen for all the artificially created short MBNL isoforms, with the partial exception of opossum MBNL3, which, although to a lesser extent than its long isoform, maintained a GCU preference (Fig. 2f-j). These results are in agreement with previous work suggesting the binding preferences of canonical MBNL proteins are largely driven by ZF2 ^32,42,43^, and suggest that the changed binding properties of the full-length mouse MBNL3 isoform may be due to a reduction in ZF2 binding dominance likely driven by the eutherian-specific mutations we identified (Extended Data Fig. 3a).

To investigate the molecular basis of the changes in binding preferences seen for the long isoform of mouse MBNL3, we next generated a number of chimeric Mouse-Opossum proteins. Introducing the two opossum zinc-finger pairs into mouse MBNL3 resulted in the mouse protein reverting to an ancestral like binding preference (Fig. 2g-j and Extended Data Fig. 3c,d). However, in contrast, introducing the mouse zinc-finger pairs into opossum MBNL3 only resulted in a modest change in binding preference. Specifically, although the consensus sequence remained strongly GCU-biased (Fig. 2i and Extended Data Fig. 3c), motifs containing a GCA sequence contributed to 20.0% of the accumulated Z-score of the top 100 motifs, while this contribution was only 8.1% for the wild type opossum MBNL3 protein (Extended Data Fig. 3d). Given that the linker region may also play a role in RNA binding ^25,28,29^, we made an additional chimeric protein consisting of the opossum protein with the mouse linker. Similar to the opossum MBNL3 with the mouse zinc-fingers, this chimeric protein showed a modest in GCA preference at the expense of GCU when compared to the wild type protein (21.7% vs. 8.1%; Fig. 2i and Extended Data Fig. 3d). Together, these data suggest that the novel binding preference of eutherian MBNL3 is a result of sequence changes in both the zinc finger and linker regions.

### Mbnl3 *restricts placental growth in mouse development*

To explore novel and ancestral physiological roles of *Mbnl3* in the developing placenta, we then made use of a mouse line in which both long and short *Mbnl3* isoforms had been knocked out ^44^. In addition to analyzing *Mbnl3* single knockouts (KO), we also examined *Mbnl2* knockouts ^32^ and the effect of *Mbnl2/3* double knockout (DKO), since, despite the divergence of *Mbnl3* in eutherians, both proteins share 64% of sequence identity in mouse and are expressed in trophoblastic tissues. Strikingly, *Mbnl3* KO resulted in a significant increase in placental weight at embryonic day 13.5 (E13.5) and E18.5 (Fig. 3a and Extended Data Fig. 4a; *P* < 0.001, Wilcoxon rank-sum tests). Analysis of placentas harvested at earlier stages revealed this weight increase first becomes apparent at E11.5 (Extended Data Fig. 4b). In contrast, no significant changes in placenta weight were seen in DKOs, whilst placentas from *Mbnl2* KOs were found to be slightly smaller than those from their wild type littermates at E13.5 (Fig. 3a; *P* = 0.036, Wilcoxon rank-sum test). To further characterize the effects of the KOs on placental development, haematoxylin and eosin staining was performed at E18.5 (Fig. 3b) and a number of phenotypic abnormalities of varying severity were observed. In DKOs, these included placental disorganization, with substantial quantities of junctional tissue (spongiotrophoblast and trophoblast glycogen cells) found within the labyrinth zone (Fig. 3c), and an increased amount of eosinophilic necrotic tissue within the junctional zone and decidua (Fig. 3d). Similar, but much less severe histological abnormalities were observed in *Mbnl2* and *Mbnl3* single KOs (Fig. 3b-d). Furthermore, an increase in the amount of both labyrinth and junctional zone tissue was seen in *Mbnl3* KO placentas, in line with their increased placental weight (Extended Data Fig. 4c-e).

**Fig. 3.**
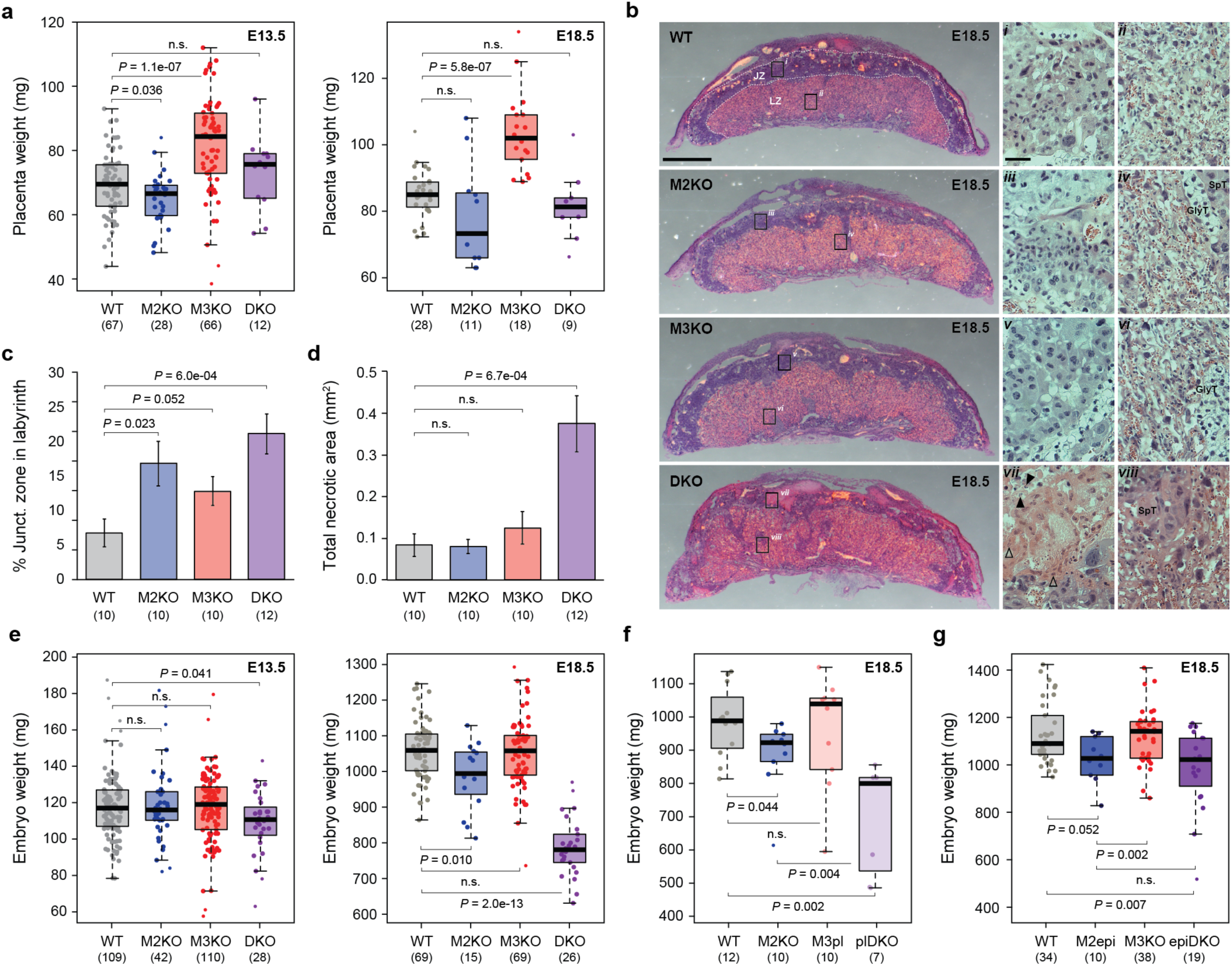
*Mbnl3* restricts placental growth. **a**,**e**, Analysis of the effects of *Mbnl2* knockout (M2KO), *Mbnl3* knockout (M3KO) or *Mbnl2*:*Mbnl3* double knockout (DKO) on placental (**a**) and embryo (**e**) weight at E13.5 and E18.5. **b**, Haematoxylin and eosin staining of WT and KO placenta sections obtained at E18.5. In the zoomed images note the glycogen trophoblast (GlyT) and spongiotrophoblast (SpT) cells within the labyrinth zone of the KO placentas (iv, vi, viii). Also note the nuclei undergoing karyorrhexis (black arrow head) and karyolysis (empty arrow head) in the DKO junctional zone (vii). JZ, junctional zone. LZ, labyrinth zone. Scale bar for full placental sections=1mm. Scale bar for zoomed images=100µm. **c**,**d**, Quantification of the proportion of total junctional zone tissue found within the labyrinth (c) and total area of esophilic necrotic tissue (d) at E18.5. A single medial section was analysed per placenta. **f**, Analysis of the effects on embryo weight of placenta specific *Mbnl3* knockout with (plDKO) or without (M3pl) universal *Mbnl2* knockout. **g**, Analysis of the effects on embryo weight of epiblast/fetus specific *Mbnl2* knockout with (epiDKO) or without (M2epi) universal *Mbnl3* knockout. For (**a**) and (**c-g**) The number of placentas/embryos analysed for each genotype are indicated in brackets and significance levels are calculated by Wilcoxon rank-sum tests.

To assess how these differences in placenta development may affect placenta function and thus embryo development, we next analyzed embryo size. *Mbnl3* KO alone had no effect on embryo weight, even at late gestational stages (Fig. 3e). In contrast, DKO embryos suffered from intrauterine growth restriction; this was severe at E18.5 (*P* < 0.001, Wilcoxon rank-sum tests) but could be observed from as early as E13.5 (Fig. 3e; *P* = 0.041, Wilcoxon rank-sum tests). A modest but significant growth restriction was also seen in *Mbnl2* single KO embryos at E18.5 (Fig. 3e; *P* = 0.010, Wilcoxon rank-sum tests). To test if the decreased embryo size seen in the DKOs was a direct result of loss of placental and not embryonic *Mbnl2/3*, we used two conditional KO strategies. Firstly, we made use of the X-linked nature of *Mbnl3*, which means that placental expression occurs almost exclusively from the maternal allele in mouse ^8^. Therefore, female embryos in which the maternal allele is KO and the paternal allele is wild type are effectively KO for *Mbnl3* in the placenta and have mosaic *Mbnl3* expression in embryonic tissues (Extended Data Fig. 4f). *Mbnl2* KO embryos with placenta-specific *Mbnl3* KO showed a similar growth restriction to full DKO embryos (Fig. 3f). Secondly, we made use of a paternal *Sox2*-Cre mouse line ^45^ to specifically knock out *Mbnl2* from the epiblast (Extended Data Fig. 4g). This resulted in a similar size reduction to that seen in full *Mbnl2* KOs, suggesting this is due to a role for this gene in the embryo not the placenta (Fig. 3g). *Mbnl3* KO did not accentuate this phenotype. Therefore, data from conditional KOs show that the stronger embryo growth restriction seen in full DKOs compared to single *Mbnl2* KOs is specifically due to a role for *Mbnl2* and *Mbnl3* in placenta. Altogether, these results indicate that both genes act partly redundantly in this tissue to ensure proper placental development, and that *Mbnl3* exerts unique functions whose disruption leads to increased placenta size.

### Mbnl2 *and* Mbnl3 *co-regulate alternative splicing and polyadenylation in placenta*

To investigate the molecular bases of these phenotypes as well as to gain insight into the mechanism of action of *Mbnl2* and *Mbnl3* in placenta, we next performed RNA-seq of placental samples harvested at E13.5 and E18.5. MBNL proteins have been extensively characterized as regulators of alternative splicing; therefore, we began by using this data to assess the effect of each KO on exon inclusion. At both time points substantially more and larger splicing changes were seen in DKO than single KO placentas (Fig. 4a,b and Extended Data Fig. 5a), indicating that *Mbnl2* and *Mbnl3* regulate an overlapping set of events and partially compensate for one another’s loss. In agreement with previous reports ^33,41,46^, the number and extent of splicing changes in *Mbnl2* KOs were greater than those in *Mbnl3* KOs (Fig. 4a-c and Extended Data Fig. 5a), and no evidence for substantial *Mbnl3*-specific splicing effects was observed. We also found no evidence of *Mbnl3* acting in placenta as a competitive inhibitor of splicing regulation by *Mbnl2* ^40,44^, as almost all splicing events significantly altered in DKOs were misregulated in the same direction in *Mbnl2* and *Mbnl3* single KOs (Fig. 4c). In line with the known function of MBNL proteins in regulating differentiation-associated alternative splicing ^47^, we found that misregulated exons acquired more immature-like inclusion levels, as shown both by analyzing splicing changes occurring between E13.5 and E18.5 in wild type placentas (Fig. 4d) and between embryonic stem cells and differentiated tissues (Extended Data Fig. 5b). Functional enrichment analysis for genes containing mis-regulated exons identified functions related to cell signaling, migration and adhesion that could relate to the disrupted placental morphology seen in the DKOs (Supplementary Tables 7 and 8). Among the most mis-regulated events were alternative exons in *Numb* (Fig. 4e), *Gpr137* (Fig. 4f) and *Numa1* (Extended Data Fig. 5c), whose immature-like inclusion patterns have been shown to inhibit differentiation, decrease epithelial tightness and associate with increased cell proliferation, respectively ^48-51^.

**Fig. 4.**
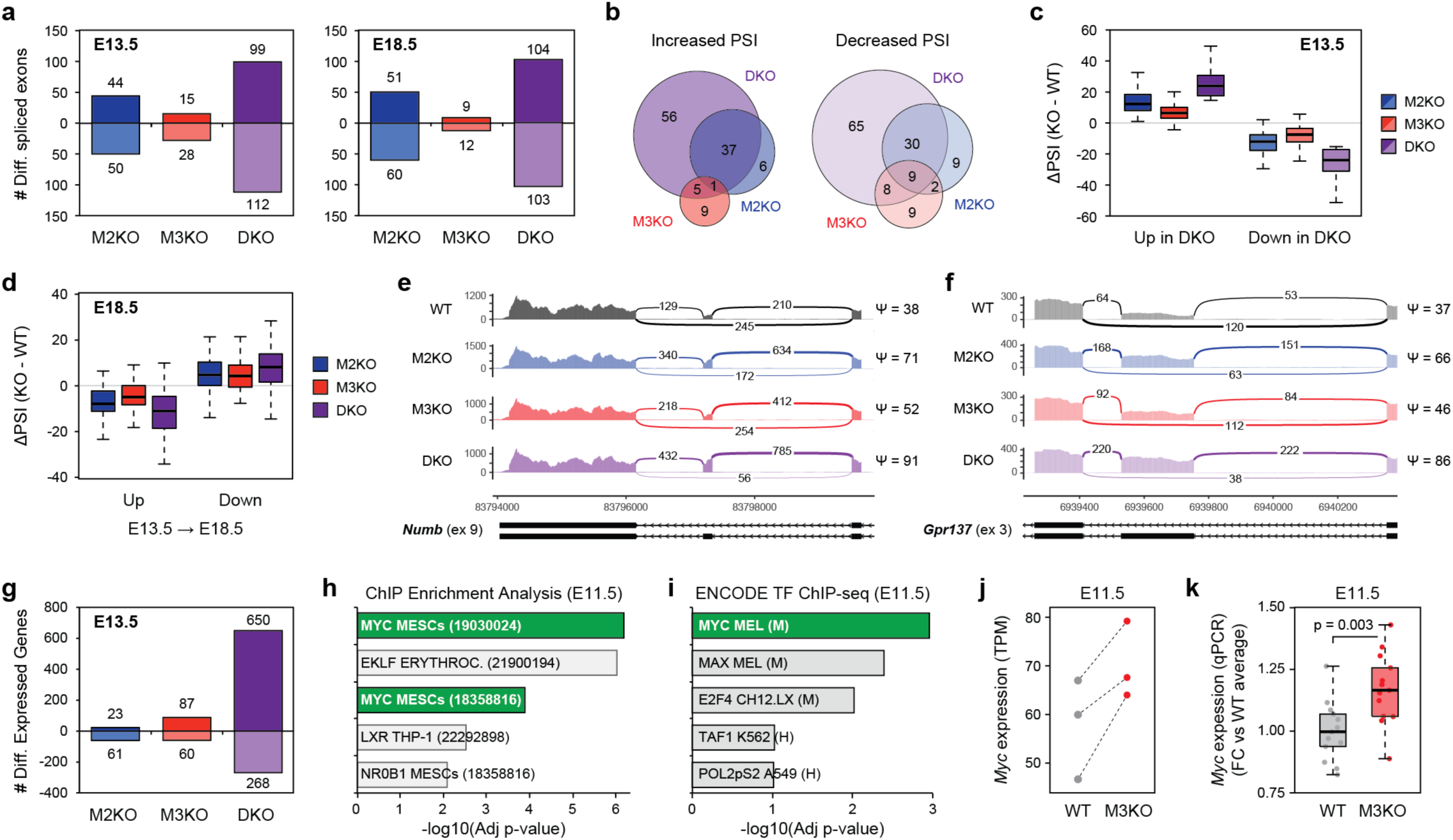
Effect of *Mbnl2, Mbnl3* and double *Mbnl2/3* knockout on the placental transcriptome. **a**, Bar charts showing the number of differentially spliced exons in *Mbnl2* KO (M2KO), *Mbnl3* KO (M3KO) and double KO (DKO) placentas at the indicated embryonic stages. Exons with increased inclusion in KOs are tallied above the x-axis whilst exons with reduced inclusion are tallied below. **b**, Venn diagrams showing the overlap in differential spliced exons between M2KO, M3KO and DKO placentas at E13.5. **c**, Boxplots showing the difference in exon inclusion (percentage spliced in, PSI) in the indicated KOs vs WT placentas for all exons differentially spliced in DKOs vs WTs at E13.5. **d**, Boxplots showing the change in PSI in the indicated KO vs WT placentas at E18.5 for all exons with differential inclusion between E13.5 and E18.5 WT placenta. **e**,**f**, Sashimi plots showing usage of *Numb* exon 9 (e) and *Gpr137* exon 3 (f) in placentas with the indicated genotypes at E13.5. The PSI values for the exons in the different genotypes are shown to the right of the plots. **g**, Bar charts showing the number of differentially expressed genes in M2KO, M3KO and DKO placentas at E13.5. Genes with increased expression in KOs are tallied above the x-axis whilst genes with reduced expression are tallied below. **h**,**i** Bar charts showing significance levels for enrichment of transcription factor (TF) binding sites amongst genes up-regulated in *Mbnl3* KO placentas at E11.5. The top 5 most significantly enriched TFs are shown for ChEA (h) and ENCODE (i) data sets. **j**,**k** Plots showing *Myc* expression levels in WT and *Mbnl3* KO placentas at E11.5 based on RNA-sequencing (j) and qPCR (k). In (j) matched placenta pools derived from the same litters are indicated by dashed lines. For (k), values correspond to the fold change between each tested placenta and the average of WT placenta; N = 13 placentas for each genotype. Significance levels were calculated by Wilcoxon rank-sum tests.

In addition to alternative splicing regulation, MBNL proteins have also been reported to modulate alternative polyadenylation site usage and MBNL3, in particular, has been found to preferentially bind the 3’
s end of transcripts ^52^. Therefore, we next generated 3’ sequencing data from the same set of placenta samples. As with alternative splicing, we found MBNL2 and MBNL3 to redundantly regulate polyadenylation site usage, and their loss to lead to more immature-like usage patterns (Extended Data Fig. 6a-d). However, in contrast to alternative splicing, relatively few genes showed significant alterations in polyadenylation site usage at E13.5 (Extended Data Fig. 6a). Among the mis-regulated targets (Supplementary Table 9) was the transcriptional repressor *Mbd2*, which showed an increase in the use of a proximal polyadenylation site from ∼0% of transcripts in the wild types to ∼50% in the DKOs (Extended Data Fig. 6e). Use of this site results in the production of a shortened protein isoform that promotes pluripotency in embryonic stem cells, whereas the long isoform enhances cell differentiation ^53,54^. This result suggests a similar mechanism likely operates in trophectodermal tissue.

### Misregulated gene expression patterns confirm maturation defects in DKO placentas

To further characterize the molecular changes occurring in single and double knockout placentas, we next used the RNA-seq data to examine differences in steady-state mRNA levels (hereafter gene expression). Overall, the results from this analysis mirrored those from splicing and polyadenylation site usage, with substantial changes in gene expression being seen in DKOs and only moderate changes in *Mbnl2* and *Mbnl3* single KOs (Fig. 4g and Extended Data Fig. 7a-c). Maturation defects were also observed at the level of gene expression in E18.5 KO placentas (Extended Data Fig. 7d). Indeed, amongst the genes up-regulated in DKO placentas at this stage were a number of transcription factors associated with trophoblast stem cell maintenance, including *Esrrb* ^55^, *Eomes* ^56^ and *Cdx2* ^57^ (Supplementary Table 10). This was accompanied by a reduction in expression of a number of genes involved in labyrinth formation and maturation, including the transcriptional regulators *Gcm1* ^58^ and *Cebpa* ^59^, and the *Gcm1* target *SynB*, which plays a role in labyrinth cell fusion ^60,61^ (Supplementary Table 10). Gene ontology analyses of differentially expressed genes revealed multiple functions altered in DKOs, including terms related to cell adhesion, migration and extracellular matrix organization (Supplementary Table 11).

### Mbnl3 *loss leads to up-regulation of Myc and its targets at E11*.*5*

In contrast to alternative splicing and polyadenylation usage, analysis of gene expression at E13.5 revealed a subset of genes that appeared to robustly respond to *Mbnl3* but not *Mbnl2* loss (Extended Data Fig. 7b,e). These genes were not significantly misregulated in *Mbnl2* KOs and had similar levels of mis-expression in *Mbnl3* KOs and DKOs (Extended Data Fig. 7e and Supplementary Table 10). This indicates that eutherian *Mbnl3* has some regulatory activities not possessed by *Mbnl2*, in line with its unique morphological phenotype. Consistent with the X-linked nature of *Mbnl3* and the expected resulting maternal bias as well as the increase in placenta size, amongst these genes were a number with a potential role in nutrient uptake, *Slc2a3, LipG, Atp1b1, Steap3* and *Slc19a2*, which where all up-regulated following *Mbnl3* loss (Supplementary Table 10).

To further investigate the molecular bases behind the increase in placenta growth specifically seen in *Mbnl3* KOs, we additionally performed RNA-seq of *Mbnl3* KO placentas at E11.5. This is the stage when the increase in placental size was first observed (Extended Data Fig. 4b), and a time of proliferative growth for the organ ^62,63^. Differential gene expression analysis at this stage revealed 10 genes to be significantly up-regulated and 23 genes to be significantly down-regulated, over half of which came from a single genomic cluster and were members of the carcinoembryonic antigen (Cea) gene family (Supplementary Table 10). Extensive examination of the genes showing significantly altered expression did not reveal any obvious candidates that may be responsible for the observed increase in placenta size. Therefore, we next performed a series of functional and regulatory enrichment analyses using looser cut-offs for differential gene expression (padj < 0.1, log2FC > 0.1). These revealed enrichment for GO terms consistent with increased growth amongst up-regulated genes, including ribosome, translation and metabolic processes (Supplementary Table 12). In addition, they showed that the pro-proliferative transcription factor *Myc* has the most significant level of binding site enrichment in the up-regulated genes according to both ChEA and ENCODE ChIP datasets (Fig. 4h,i and Supplementary Table 12). Similarly, *MycN* was found to be the top candidate in a transcription factor perturbation dataset (Supplementary Table 12). In line with these results, expression of *Myc* was up-regulated ∼1.2 fold at E11.5, both based on RNA-seq data and qPCR assays using independent placenta samples (Fig. 4j,k). Up-regulation of *Myc* and its associated regulatory network is consistent with an increase in proliferation contributing to the larger placentas seen in *Mbnl3* KO mice.

### Mbnl3 *KO alleviates fetal growth reduction following maternal calorie restriction*

The increase in placental growth seen on *Mbnl3* KO is consistent with the theory that the eutherian X-chromosome is enriched for placentally expressed genes that favor increased maternal fitness ^9-11^. In this context, *Mbnl3* would act by limiting the amount of resources allocated to placenta growth, and thus fetal development. However, whilst there is an obvious maternal cost in terms of energy expenditure for increasing placenta size in *Mbnl3* KO animals, we did not observe any fetal advantage, such as increased embryo size. This could be either due to a suboptimal placenta function in *Mbnl3* KOs or because embryo size is already optimal under standard laboratory conditions, where mothers have *ad libitum* access to food. The second possibility led us to hypothesize that the increased placental size seen in *Mbnl3* KOs may provide additional placental reserve capacity to *Mbnl3* KO embryos when resources are limited. To test this, we examined the effects of *Mbnl3* KO on fetal growth following mid-gestational maternal calorie restriction, which results in a reduction in embryo size of ∼50% at E18.5 (Extended Data Fig. 8a; ^64^). Under these conditions, placentas from *Mbnl3* KOs were still found to be larger than those of their wild type littermates (Fig. 5a,b and Extended Data Fig. 8b). However, in contrast to non-caloric restricted conditions and in agreement with our hypothesis, analysis of embryo weights revealed *Mbnl3* KO fetuses from calorie restricted pregnancies to be significantly larger than their wild type littermates at E18.5 (Fig. 5c,d and Extended Data Fig. 8c).

**Fig. 5.**
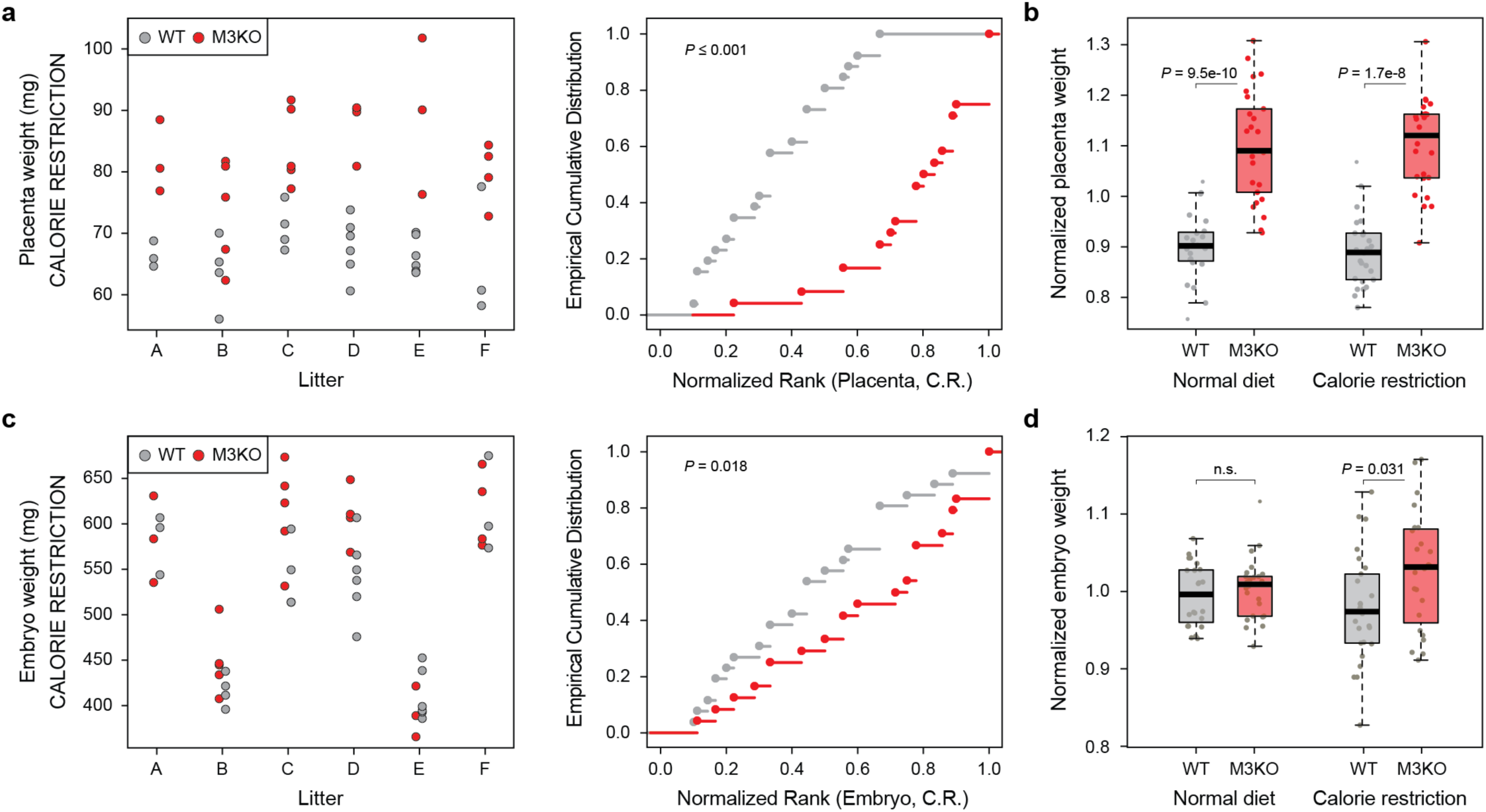
*Mbnl3* KO reduces the effect of mid-gestational maternal calorie restriction on fetal growth. **a**,**c** Dot plots showing the weights of individual WT (grey) and *Mbnl3* KO (red, M3KO) placentas (a) and embryos (c) harvested from six calorie restricted mothers (A-F) and the corresponding empirical cumulative distribution plot for each genotype. Significance levels were calculated by a permutation test with 1000 iterations swapping the genotype labels within each litter. **b**,**d**, Analysis of the effects of maternal calorie restriction on the weight of WT and M3KO placentas (b) and embryos (d) at E18.5. For each embryo/placenta, the weight was normalized with respect to the average of WT and KO weights within each litter. Significance levels are calculated by Wilcoxon rank-sum tests.

## Discussion

We have identified two related RNA binding proteins, MBNL2 and MBNL3, with differing evolutionary histories that play both overlapping and distinct roles in placenta development. Despite its accelerated sequence evolution, *Mbnl3* appears to have retained sufficient ancestral regulatory activity to act partially redundantly with *Mbnl2* to control placental morphology and physiology and thus facilitate proper fetal growth. In particular, *Mbnl2* and *Mbnl3* appear to be required for proper placenta transcriptomic maturation, organization of the labyrinth and junctional zones, and to prevent excessive cell death/necrosis at the maternal fetal interface. The need for MBNL proteins for transcriptomic maturation in placenta is in line with that observed in other tissues and likely occurs via both co- and post-transcriptional regulation of mRNA metabolism, particularly alternative splicing ^32,47,52,53,65-68^.

On the other hand, *Mbnl3* has acquired a new biological function limiting placental growth. The increase in placental size in mouse *Mbnl3* KOs is first seen at around E11.5. This is a time when placental growth is largely driven by cell proliferation and not an increase in cell size, which occurs from E12.5 onwards ^62,63^. Furthermore, the pro-proliferative transcription factor *Myc* is upregulated in *Mbnl3* KOs at E11.5 and genes upregulated in these KOs are enriched for *Myc* binding within their promoters. Altogether, this suggests that *Mbnl3* limits placental growth at least in part by reducing cell proliferation in association with lower levels of *Myc* expression. The effects of *Mbnl3* on *Myc* levels are unlikely to be direct, as previous CLIP-seq experiments in other cell types with robust *Myc* expression have found no evidence that MBNL3 binds directly to *Myc* transcripts ^41,52^. Therefore, it remains unclear how *Mbnl3* exerts its effects on placental growth; however, it is tempting to speculate that it relates to the evolution of its novel RNA binding specificity and/or of a eutherian-specific short isoform with a single pair of zinc fingers (see Supplementary Discussion for details). Previous studies have found the two *Mbnl3* isoforms to be in distinct cytoplasmic compartments, with the long isoform largely localized to ribosomes and the short isoform largely soluble, suggesting that they play distinct roles in RNA metabolism ^41^. Given that both isoforms are strongly expressed in mouse placenta and have similar binding preferences that are distinct from those of canonical MBNL proteins, they could modulate different aspects of the metabolism of the same RNA transcripts in a coordinated manner to inhibit growth. However, the short isoform appears to be more abundant in both cow and humans, hinting that it may harbor the majority of this activity. Irrespectively, the important regulatory interactions likely occur in the cytoplasm as eutherian *Mbnl3* has lost an alternative exon driving nuclear localization and has a more cytoplasmic localization, a higher ratio of 3′ UTR to intronic binding, and is a weaker splicing regulator than other MBNL proteins (this paper and ^33,41,52^).

Irrespectively of the molecular mechanisms of action, the evolution of *Mbnl3*’s function to restrict placental growth is likely linked to its genomic location on the X chromosome. In line with this, whereas both *Mbnl2* and *Mbnl3* appear to have been recruited to the placenta early in eutherian evolution, this recruitment has been accompanied by loss of expression from other tissues and specialization only in *Mbnl3*, but not in *Mbnl2*. Due to paternal-specific inactivation in placenta, the mouse X-chromosome is expected to accumulate alleles that favor the maternal side of the parental conflict ^9-11^. The maternal cost of a larger placenta is an increase in energy requirements, whilst we demonstrate here one of the paternal advantages is better maintenance of fetal growth following maternal calorie restriction, an event that could conceivably be a frequent occurrence in an animal’s natural environment. Altogether, this suggests an evolutionary model where changes to the *Mbnl3* loci leading to reduced placental growth have been favored due to its genomic location and have either driven and/or been facilitated by its tissue-restricted expression, consolidating *Mbnl3* as a novel player in the parental genetic conflict in eutherians.

## Methods

### Identification of splicing factors enriched in trophoblastic tissues

We used a list of previously defined RNA binding proteins and other genes known to be involved in splicing and/or alternative splicing regulation ^69,70^. We restricted the analysis to 197 splicing factors with one-to-one orthologs between mouse and human based on Ensembl Biomart. One-to-one orthologs vs mouse were also retrieved for cow and opossum, when available. All studied splicing factors for the four species are listed in Supplementary Table 13.

To investigate the enrichment of these splicing factors in trophoblastic tissues in mouse, we focused on three comparisons: (i) placenta vs. non-placental tissues, (ii) trophoblast vs. inner cell mass at the blastocyst stage, and (iii) trophoblast stem cells vs. embryonic and extraembryonic endoderm stem cells. Gene expression quantifications, using the Transcript Per Million (TPM) metric, for public RNA-seq data for different placenta and non-placenta tissues as well as embryonic blastomeres and embryonic stem cells were obtained from *VastDB* ^71^, assembly mm10. SRA identifiers and associated information are listed in Supplementary Table 1. In addition, for comparison (iii) we generated *de novo* RNA-seq of different lines of cultured trophoblast stem (TS) cells and extraembryonic endoderm stem (XEN) cells. Culture details and sequencing and mapping information are provided below and in Supplementary Table 1. TPM values for TS and XEN samples were obtained using *vast-tools* v2.5.1 ^71^.

For comparison (i), for each gene we first calculated the average TPM expression for each tissue type (groupings are provided in the column “Group” of Supplementary Table 1). Next, we calculated the log2 fold change of expression between placental tissues and the average of non-placental tissues, which are plotted in Fig. 1a and shown in Supplementary Table 2. Only genes with TPM > 10 in placental tissues were considered. For the other two comparisons (Fig. 1b and c), we followed a similar logic and averaged the expression of each gene in (ii) trophectoderm (TE) and (ICM) blastomeres at E3.5 and (iii) trophoblast stem cells (TS) and the average of embryonic stem cells and extraembryonic endoderm stem cells (ESC&XEN).

To compare placental and non-placental tissues in human, cow and opossum we used a similar approach to the one described for mouse, relying on public RNA-seq data. For human (hg38) and cow (bosTau6), we retrieved TPM values from VastDB ^71^. For opossum and non-mammalian vertebrates, we collected RNA-seq data from the Short Read Archive (SRA) (Supplementary Table 1) and calculated TPMs using *vast-tools* v2.5.1 for each species.

### Analysis of Mbnl3 transcription start sites and alternative splicing

To identify potential transcription start sites (TSS) and alternative splicing events leading to different N-terminal isoforms of MBNL3, we looked for competing splicing donors (5′ splice sites) that are spliced to the splicing acceptor (3′ splice site) of the second coding exon, which contains the in-frame ATG giving rise to the short protein isoform and that is present in all known *Mbnl3* transcripts. For this purpose, we took advantage of *vast-tools align*, which generates an output file for each RNA-seq sample (sample.eej2) containing the read counts for all exon-exon junctions annotated in the *vast-tools* junction-based library ^71^. The reference splicing acceptor for *Mbnl3* for each species were: 11 from ENSMUSG00000036109 (mm10), 16 from ENSG00000076770 (hg38) and 0 from ENSBTAG00000014088 (bosTau6). The corresponding donors are numbered starting from 0 (the most upstream) for each species, and the coordinate on the X-chromosome and its location in the genome are provided in Extended Data Fig. 2a-c and Supplementary Table 6. Each donor was grouped and colored based on the evolutionary origin of the associated TSS region and whether or not it would lead to a short MBNL3 protein isoform: black, long isoform from the ancestral TSS; grey, short isoform from the ancestral TSS by skipping of the first coding exon; red, short isoform from the eutherian-specific TSS; orange, short isoform from the rodent/primate-specific TSSs. To provide the percent of reads coming from each group of donors (Fig. 2c and Extended Data Fig. 2e,f), we calculated the percent of reads for each of those groups out of the total number of exon-exon junction reads to the reference acceptor as provided in the sample.eej2 files and, then, we grouped similar samples (e.g. neural tissues, muscle, etc.) as indicated in the column “GROUP” of Supplementary Table 6.

### Western blotting

Western blot analysis was conducted according to standard laboratory procedures. Lysates were prepared from staged placental samples by disruption with a sterile pestle in Laemmli buffer (0.05M Tris-HCl at pH 6.8, 1% SDS, 10% glycerol, 0.1% b-mercaptoethanol) on ice. Samples were then boiled at 95°C for 10 min, sonicated using a bioruptor (10cycles 30 secs on, 30 secs off), and spun at 20,000g for 90 secs. Proteins were separated by 4-15% SDS-PAGE gel (Bis-Tris Criterion™ TGX™ pre-cast gels, Bio-Rad) and transferred to PVDF membranes (Thermo Scientific). Membranes were incubated overnight with MBNL3 antisera ^41^ at 1:500 or anti Lamin B1 (1:1000, Cell Signaling Technologies) in 5% non-fat dairy milk in PBS plus Igepal (0.05%, Merck).

### Mouse transgenic reported assays

Three fragments of the putative enhancer/promoter region surrounding the eutherian specific transcriptional start site of *Mbnl3* were amplified from mouse genomic DNA using the primers shown in Supplementary Table 14. Fragments were then cloned together into a modified pBluescript vector ^72^ containing a lacZ reporter gene. Prior to microinjection the construct was linearized and plasmid sequences removed. The construct was then microinjected at 3–6 ng/µl into the pronucleus of fertilized mouse oocytes at E0.5. Embryos were collected from E6.5 to E7.5, fixed, and stained for β-galactosidase activity.

### Molecular evolution analyses for Mbnl2 and Mbnl3

Protein sequence alignments and tree production were made using the CLC main work bench (ver. 5.5.1, Qiagen) software package. MBNL isoforms used for alignment only contained the coding exons common to all *Mbnl2* and *Mbnl3* orthologs (coding exons 5 [54bp] and 9 [∼74bp] were excluded from all aligned sequences). Deeply conserved residues (red box) were defined as those conserved across MBNL2 and non-eutherian MBNL3 proteins whereas all other residues were considered non-deeply conserved (green boxes). The protein tree was constructed by the neighbor Joining method with Kimura protein corrections. To investigate loss of coding exon 5 (54bp), *Mbnl2* and *Mbnl3* orthologs were first examined on the Ensembl (version 102) and UCSC (http://genome.ucsc.edu) genome browsers. For *Mbnl* genes where exon 5 was not annotated exon loss was confirmed by manual inspection of the intronic region.

### MBNL constructs used for RNAcompete

For all *Mbnl* constructs tested the exon usage pattern followed that of the predominant MBNL3 isoform in mouse placenta: all MBNL proteins tested included alternative coding exon 8 (95/98bp) and lacked alternative coding exons 5 (54bp), 7 (36bp) and 9 (∼74bp). Mouse *Mbnl* isoforms were cloned from placenta/limb cDNA using the primers detailed in Supplementary Table 14. DNA sequences corresponding to opossum and chicken MBNL isoforms alongside the chimeric MBNLs were synthesized *in vitro* by Bio Basic and IDT, respectively. Corresponding short isoforms were generated either from genomic DNA or cloned long isoforms using the primers detailed in Supplementary Table 14.

### RNAcompete and associated bioinformatic analyses

The RNA pool generation, RNAcompete pull-down assays, and microarray hybridizations were performed as previously described ^34-36^. Briefly, GST-tagged RNA-binding proteins (20 pmoles) and RNA pool (1.5 nmoles) were incubated in 1 mL of Binding Buffer (20 mM Hepes pH 7.8, 80 mM KCl, 20 mM NaCl, 10% glycerol, 2 mM DTT, 0.1 mg/mL BSA) containing 20 mL glutathione sepharose 4B (GE Healthcare) beads (pre-washed 3 times in Binding Buffer) for 30 minutes at 4°C, and subsequently washed four times for two minutes with Binding Buffer at 4°C.

One-sided Z-scores were calculated for all 7-mer motifs as described previously ^34^. Z-scores approximate the affinity of the RNA binding protein for that specific 7-mer. To obtain a measure of the relative contribution of a specific motif (e.g. GCUU or GCAGC) to the overall affinity, we summed the Z-scores of all 7-mers containing the motif among the top 100 7-mers and calculated the fraction of that sum with respect to that of all 100 7-mers (Fig. 2g). Similarly, to assess the relative contribution of the nucleotide after a GC, for each 7-mer containing a GC among the top 100 7-mers, we summed the Z-scores for 7-mers containing GCA, GCC, GCG or GCU and calculated the fraction corresponding to each of them (Fig. 2i). Finally, for those 7-mers containing GCxGC or GCxxGC among the top 100 7-mers, we calculated the relative contribution of those with GCxGC or GCxxGC motifs as the log2 ratio between the sums of their Z-scores (Fig. 2j).

### Animal husbandry, and embryo and placenta weight analysis

All protocols were carried out in accordance to the European Community Council Directive 2010/63/EU and approved by the local Ethics Committee for Animal Experiments (Comitè Ètic d’Experimentació Animal-Parc de Recerca Biomèdica de Barcelona, CEEA-PRBB, CEEA 9086 and MDS 0035P2). *Mbnl3*^*tm2*.*1Sws/+* 44^, *Mbnl2*^*tm1*.*1Sw/+* 32^, *Mbnl2*^*tm1Sws/+* 32^ and *Edil3*^*Tg(Sox2-cre)1Amc* 45^ mice alongside the required intercrosses were maintained on mixed backgrounds. Embryo genotyping was conducted on tail biopsies as described ^73^ using published primers. Embryo sex determination was based on the presence or absence of the Y-linked gene *Sry* as determined by PCR using primers detailed in Supplementary Table 14.

In all analysis WT littermates were used as controls. Furthermore, female *Mbnl3* heterozygous embryos carrying a wild type maternal allele where considered to be wild type for *Mbnl3* as it is almost exclusively expressed from the maternal allele in placenta (Extended Data Fig. 4f) and *Mbnl2* heterozygotes were considered wild type for *Mbnl2* as neither placenta nor embryo size was significantly altered in *Mbnl2* heterozygotes vs full wild type embryos regardless of the knockout status of *Mbnl3*. For the analysis of placenta specific *Mbnl3* KOs (a genotype only obtainable in female embryos) all results from male siblings of all genotypes were excluded.

For placental weight analysis at E10.5 and E11.5 only the embryonic portion of the placenta was analyzed, which was separated from the maternal decidua by blunt dissection. For placentas analyzed from E13.5 onwards the whole placenta was weighed.

### Maternal calorie restriction analysis

For maternal calorie restriction experiments pregnant dams were fed 50% of their normal daily chow intake from E11.5 until embryo harvesting ^64^. Only litters with at least three WTs and three KOs were used for the analyses. Raw values for placenta and embryo weight for each genotype within each litter are provided in Fig. 5a,c (calorie restricted diet) and Extended Data Fig. 8b,c (normal diet). To normalize for the large variability observed among litters we performed two complementary approaches. First, we plotted the values as Empirical Cumulative Distribution Function (ECDFs) to visualize the effects. To test the hypothesis that KO placentas/embryos are heavier than their littermates, we ranked each placenta/embryo within each litter and obtained a total sum of ranks per genotype within each litter. We then randomly swapped the genotype labels within each litter 1000 times and computed in how many iterations the sum of ranks was the same or smaller than in the test case. Second, we normalized each placenta/embryo measure by dividing its weight by the average of the mean WT and mean KO weights within each litter (Fig. 5b,d). Statistical significance of the difference between the normalized values was assessed using Wilcoxon rank-sum tests.

### *Whole mount* in situ *hybridization and haematoxylin and eosin staining*

Whole mount *in situ* hybridization was carried out following standard procedures ^74^. Probes were synthesized from cDNAs cloned into the pGEM®-T Easy (Promega) plasmid. The cloning primers used for probe generation are shown in Supplementary Table 14. Hematoxylin and eosin staining was conducted according to standard laboratory practice on 6µm sagittal mid-sections from placentas harvested at E18.5, fixed overnight in 4%PFA and embedded in paraffin wax. Sections from 10 or more placentas were examined per genotype.

Images of whole embryos and placenta sections were captured using an Olympus SZX16 stereo microscope and DP73 digital camera. Zoomed images were captured using a Leica DM6000 B upright microscope and DFC420 digital camera with a 20X objective. For quantification of hematoxylin and eosin stained sections, labyrinth, junctional (spongiotrophoblast and glycogen trophoblast) and neurotic tissue was manually defined based on morphology. Analysis was conducted using ImageJ (Version 1.50g, NIH, Maryland, USA). A single medial section was quantified per placenta. Total labyrinth tissue area = (total area of labyrinth zone) – (total area of junctional tissue in the labyrinth). Total junctional tissue area = (area of junctional zone) + (area of junctional tissue in the labyrinth).

### RNA sample collection for sequencing and RT-qPCR

TS^JR 75^, TS^FxL4 76^, XEN^JR 77^ and XEN^DIZ6 76^, cell lines were cultured as described ^75,77^. For E11.5 placental samples the whole embryonic portion of the placentas was separated from the maternal decidua by blunt dissection. For E13.5 and E18.5 placental samples broad medial sections of placenta were cut and tissue layers separated using forceps. For E13.5 samples the embryonic portion of the placenta was collected. For E18.5 samples only the labyrinth zone was collected to avoid signal from the necrotic cells found in the junctional zone of DKO placentas (Fig. 3b,d) overwhelming the analysis. For sequencing of E11.5 placentas 3 samples, each containing 3 placentas, were collected per genotype and samples were paired so that WTs and mutants came from the same litter(s). For sequencing of E13.5 and E18.5 placentas 2 samples, each containing tissue from a minimum of 5 placentas, were collected for each genotype. In all cases RNA was extracted using the RNeasy MiniKit (QIAGEN).

### Quantitative real time PCR

For qPCR cDNA was prepared from total RNA using Superscript III reverse transcriptase (Invitrogen) and random nonamer primers (Invitrogen). qPCR was performed using the NZYSpeedy qPCR Green Master Mix (Nzytech). Primers used are detailed in Supplementary Table 14. For analysis of *Mbnl2* and *Mbnl3* expression the house keeping genes *Ywhaz* and *Hmbs* were used for normalization. For analysis of *Myc* expression the housekeeping genes *Ywhaz, Srp14, Gapdh* and *Eef2K* were used for normalization and expression was calculated relative to average WT expression.

### Next-generation sequencing

For gene expression and alternative splicing analyses, standard polyA-selected Illumina libraries were generated using standard protocols by the CRG Genomics Unit. An average of ∼100 million (XEN and TS cells), ∼47 million (E13.5 and E18.5) 125-nt paired end or ∼47 million 50-nt single end (E11.5) reads were generated using a HiSeq2500 machine. For alternative polyadenylation analysis, the same E13.5 and E18.5 RNA samples were processed by the CRG Genomics Unit with the QuantSeq 3′ mRNA-Seq Library Prep Kit Reverse (REV) from Lexogen following manufacturer instructions. An average of ∼11 million 50-nt single end reads were generated using a HiSeq2500 machine. General and mapping statistics for all samples are provided in Supplementary Table 1.

### Alternative splicing analysis

To obtain exon inclusion levels, we processed RNA-seq samples with *vast-tools* v2.5.1 ^71^ following default options. In brief, each sample was mapped using *vast-tools align* and the mm10 VASTDB library (vastdb.mm2.23.06.20.tar.gz). Next, tables with inclusion levels using the Percent-Spliced-In (PSI) metric for all exons were generated for E13.5 and E18.5 samples separately using *vast-tools combine* with the default options. Events with sufficient read coverage across all samples within a given time point were then extracted using *vast-tools tidy* with the following options: --min_N 8 --min_SD 0 --noVLOW (i.e. only exons with LOW or higher coverage scores were allowed). From these exons, differentially spliced exons between each KO and the WT for each time point were defined as those with an average absolute change in PSI (ΔPSI) between KO and WT ≥ 15 and at least ΔPSI ≥ 5 between each of the two replicates in the same direction. Gene Ontology (GO) enrichment analyses for genes with differentially spliced exons for each KO vs WT comparison at each time point were performed using DAVID v6.8 ^78^ for GOTERM_BP_DIRECT, GOTERM_MF_DIRECT, GOTERM_CC_DIRECT and KEGG_PATHWAY. As a background set for each time point, we used genes with at least one exon event annotated in *vast-tools* with sufficient read coverage across all samples as defined above (9,211 genes for E13.5 and 8,263 for E18.5). Sashimi plots for selected events were generated using *ggsashimi* ^79^ with BAM files produced by mapping RNA-seq reads to mm10 genome using STAR ^80^.

Exons that change during placental development were defined as those with a significant change between WT samples at E13.5 and E18.5 based on the same cut-offs (|average ΔPSI| ≥ 15 and ΔPSI between each replicate ≥ 5 in the same direction). ESC-differential exons (i.e. exons that are differentially spliced between embryonic stem cells and differentiated cell and tissue types) were defined as previously reported ^47^. For this purpose, we obtained PSI values from *VastDB* for mouse (mm10 release 21/12/21) and obtained the average ΔPSI between embryonic stem cells and differentiated cell and tissue types using the script Get_Tissue_Specific_AS.pl ^81^ (https://github.com/vastdb-pastdb/pastdb) with the config file provided in Supplementary Table 15 and the option --test_tis ESC.

### Polyadenylation site usage analysis

We aligned the Lexogen Quantseq Reverse sequences to the mm11 assembly (Ensembl 104 gene annotation) using STAR with parameters --outFilterMultimapNmax 1 -- outFilterMatchNminOverLread 0.2 --outFilterScoreMinOverLread 0.2 --sjdbGTFfile. We then clustered 5’-end of alignments with inverted strand (with polyA signal filtering to avoid most internal priming events) to construct the polyA database from our data. We then used this database to compute polyA site counts (clustering of alignments around detected polyA sites with 50 down/up stream nt tolerance) and used DEXSeq ^82^ to identify regulated polyA sites in genes. We report regulated genes with polyA site pairs that undergo fold-changes in opposite directions and for which changes are significant (FDR<0.1).

For further analyses, we selected a single polyA pair per gene and stage (Supplementary Table 9). For this purpose, among the pairs (if more than one) that were significantly misregulated in the largest number of KO vs WT comparisons (from 0 to 3), we performed a ranked selection: (i) we took the pair with the largest absolute fold change if the total number of reads across the four conditions for the pair were at least 15% of the pair with the largest number of reads; (ii) else, we took the pair with the largest number of reads of all (not only significant) pairs; (iii) else, if the pair had more than 30% of the number of reads of the pair with the largest number of reads, we took the one with the highest fold change; (iv) else, we discarded the gene.

### Gene expression analysis

RNA-seq samples were used to estimate transcript abundances using *Salmon* quasi-mapping approach with default parameters plus the --seqBias and the --gcBias options to perform sequence-specific and GC bias correction respectively, and the --validateMappings option to ensure that mappings are validated using an alignment-based verification ^83^. The reference fasta file used for the quantification was the set of cDNA sequences for Ensembl and *ab initio* genes predicted in the mm10 genome assembly in Ensembl ^84^(Version 88). Gene expression estimates were obtained from transcript abundance estimates using the txtimport package ^85^ with the option countsFromAbundance=lengthScaledTPM, to obtain estimated gene counts scaled using the average transcript length over samples and the library size.

Differential gene expression analysis was performed using these gene expression estimates with the *DESeq2* package ^86^ with default settings. In WT *vs* KO differential expression analyses, genes with an FDR-adjusted p-value (padj) smaller than 0.05 (E13.5 and E18.5 samples) or 0.1 (E11.5 samples) were considered differentially expressed. For display purposes, shrunken log2 fold changes were used. To investigate the effect of the different KOs on placenta maturation, log2 fold changes for each KO at E18.5 were plotted for genes that were differentially expressed between E13.5 and E18.5 in WT samples using strict cut-offs (padj < 0.01 and |FC| > 5) (Extended Data Fig. 7d).

GO analyses for differentially expressed genes for each KO at E13.5 and E18.5 were performed using DAVID v6.8 ^78^ for GOTERM_BP_DIRECT, GOTERM_MF_DIRECT, GOTERM_CC_DIRECT and KEGG_PATHWAY. Genes quantified by *DESeq2* were provided as background set (17,488 for E13.5 and 17,221 for E18.5). For E11.5, to increase the list of potentially differentially expressed genes, we relaxed the cut-offs to select genes with padj < 0.1 and |log2FC)| > 0.1 (409 and 331 up and downregulated genes, respectively). We conducted GO enrichment analyses for these subsets of genes using DAVID as described above with all genes for which a valid p-value (non NA) is proved by DESeq2 as background set (16,972 genes). In addition, we used the gene set enrichment analyses for transcription factor (TF) binding (ChEA_2016 and ENCODE_TF_ChIP-seq_2015) and the “TF Perturbations Followed by Expression” test from the web tool *EnrichR* ^87^.

## Acknowledgements

We thank Ben Blencowe for initial support on this project and Luis P. Iñiguez for help on generating plots. The research has been funded by the Spanish Ministerio de Ciencia (BFU2017-89201-P), the ‘Centro de Excelencia Severo Ochoa 2013-2017’(SEV-2012-0208), and the People Programme (Marie Curie Actions) of the European Union’s Seventh Framework Programme (FP7/2007-2013) under REA grant agreement n° 608959 to TS. We acknowledge the support of the CERCA Programme/Generalitat de Catalunya and of the Spanish Ministry of Economy, Industry and Competitiveness (MEIC) to the EMBL partnership.

## Extended Data Figures

**Extended Data Fig. 1.**
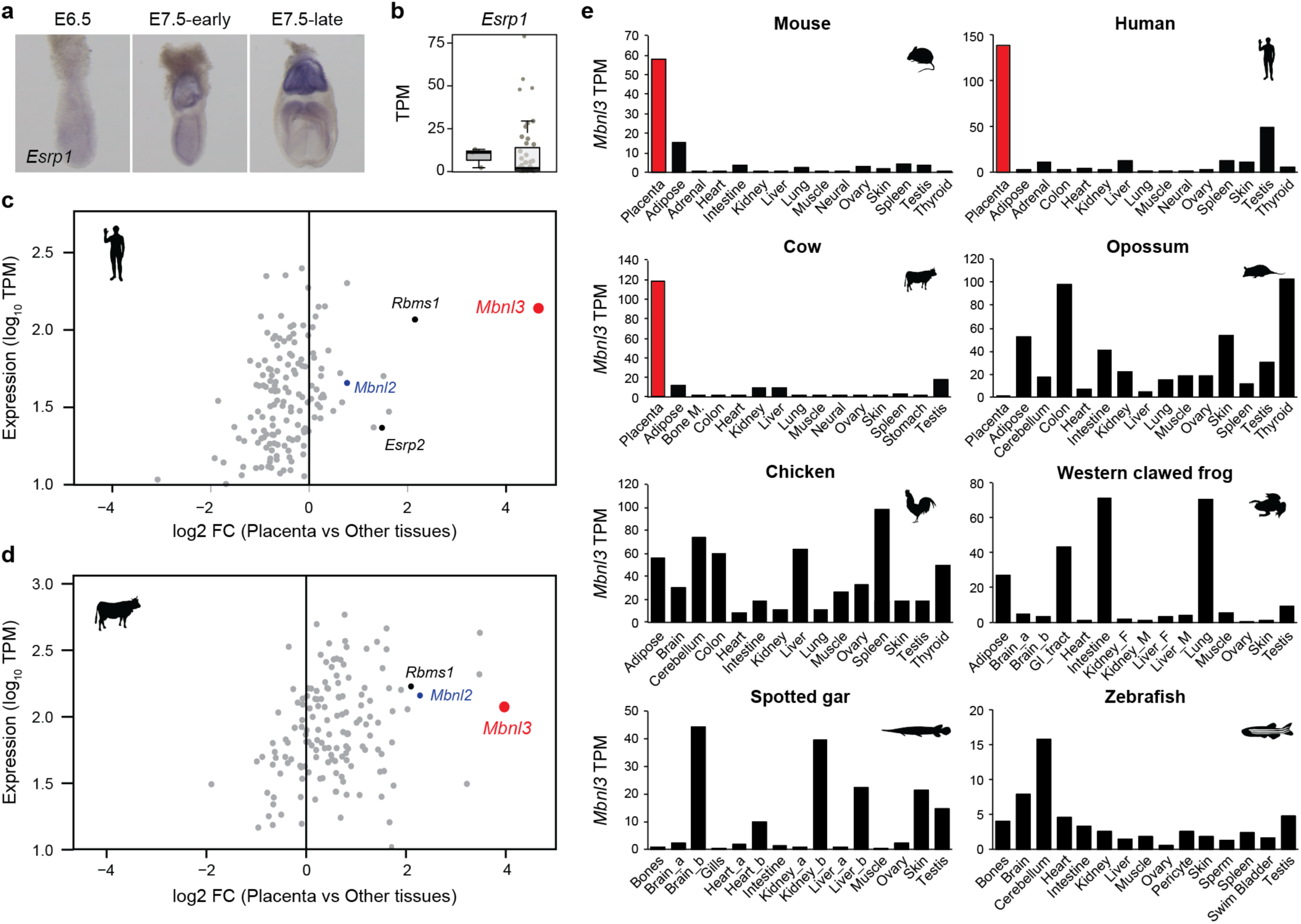
*Mbnl3* expression is highly enriched in placenta in eutherians. **a**, Whole mount *in situ* hybridisation analysis of *Esrp1* in mouse embryos at the indicated stages. **b**, Box plot showing expression levels of *Esrp1* in placenta and non-placental tissues in mouse. **c**,**d**, Scatter plots showing placental expression level and placenta vs other tissues enrichment for 197 splicing regulators in human (c) and in cow (d). **e**, Expression of *Mbnl3* orthologs from different vertebrate species across differentiated adult tissues. Placenta samples are highlighted in red. RNA-seq samples are listed in Supplementary Table 1.

**Extended Data Fig. 2.**
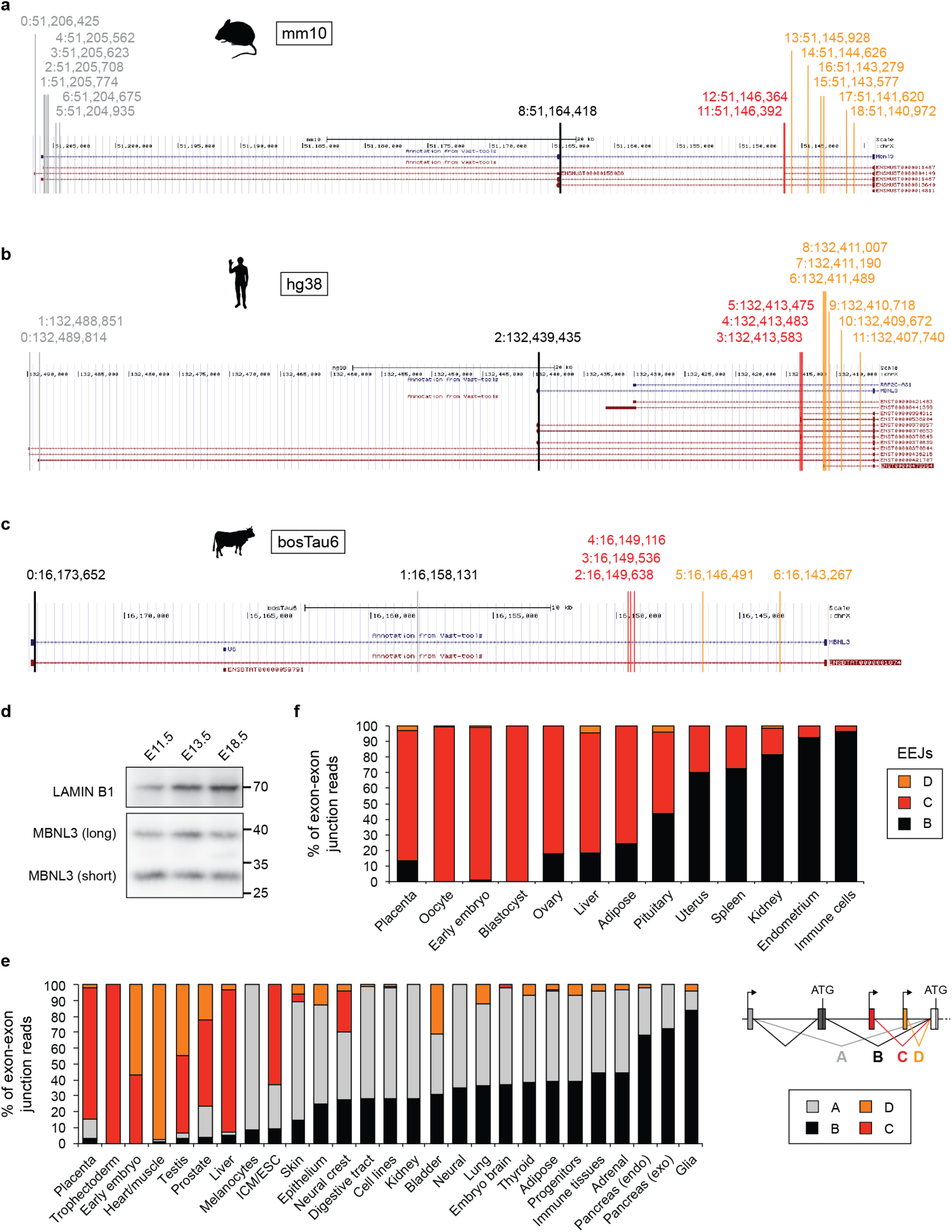
Novel transcription start sites in Mbnl3 and short isoform use. **a-c**, Genome organization for mouse (a), human (b) and cow (c) of the annotated donor splice sites upstream of the second coding exon of *Mbnl3*, which encodes the internal ATG used to translate the short isoform. The coordinate on the X-chromosome is indicated for each competing donor. The color code corresponds to that of Fig. 2a and the donor IDs correspond to that of Supplementary Table 6. **d**, Western blot showing short (∼27kDa) and long (∼38kDa) MBNL3 isoforms in placenta at different developmental time points. **e**,**f**, Percent of exon-exon junction reads from each group of competing donors (as shown in a-c and schematized in the legend) across different samples from human (e) and cow (f). Individual donor counts for each sample are listed in Supplementary Table 6.

**Extended Data Fig. 3.**
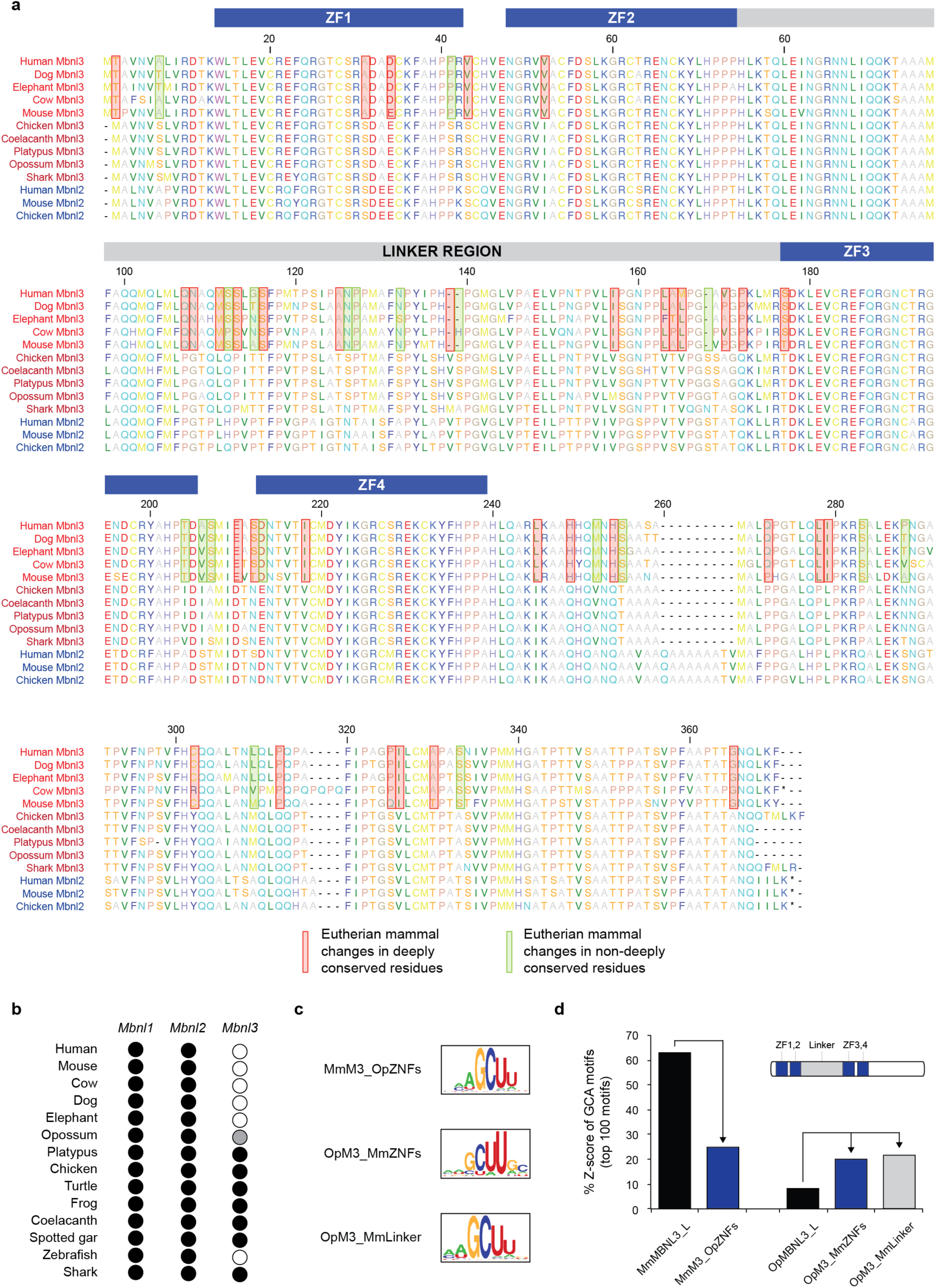
Molecular evolution of eutherian MBNL3 proteins. **a**, Sequence alignment for different MBNL3 (red) and MBNL2 (blue) proteins from various vertebrate species. Eutherian changes in deeply conserved (conserved between MBNL2 and non-eutherian MBNL3 proteins) and non-deeply conserved residues are highlighted using red and green boxes, respectively. Zinc-fingers (ZF) and the liker region are indicated. **b**, Presence (black) or absence (white) of exon 5 in different *Mbnl* genes across vertebrates. In opossum exon 5 can be identified but it contains an in-frame stop codon (grey) **c**, RNAcompete-derived sequence logos for the indicated chimeric proteins, with either mouse (Mm) or opossum (Op) zinc-finger or linker regions in the other species’ backbone. **d**, Analysis of the Z-score contribution of sequences containing GCA motifs to the total cumulative Z-score of the top 100 RNAcompete derived 7-mers for the indicated wild type and chimeric MBNL proteins.

**Extended Data Fig. 4.**
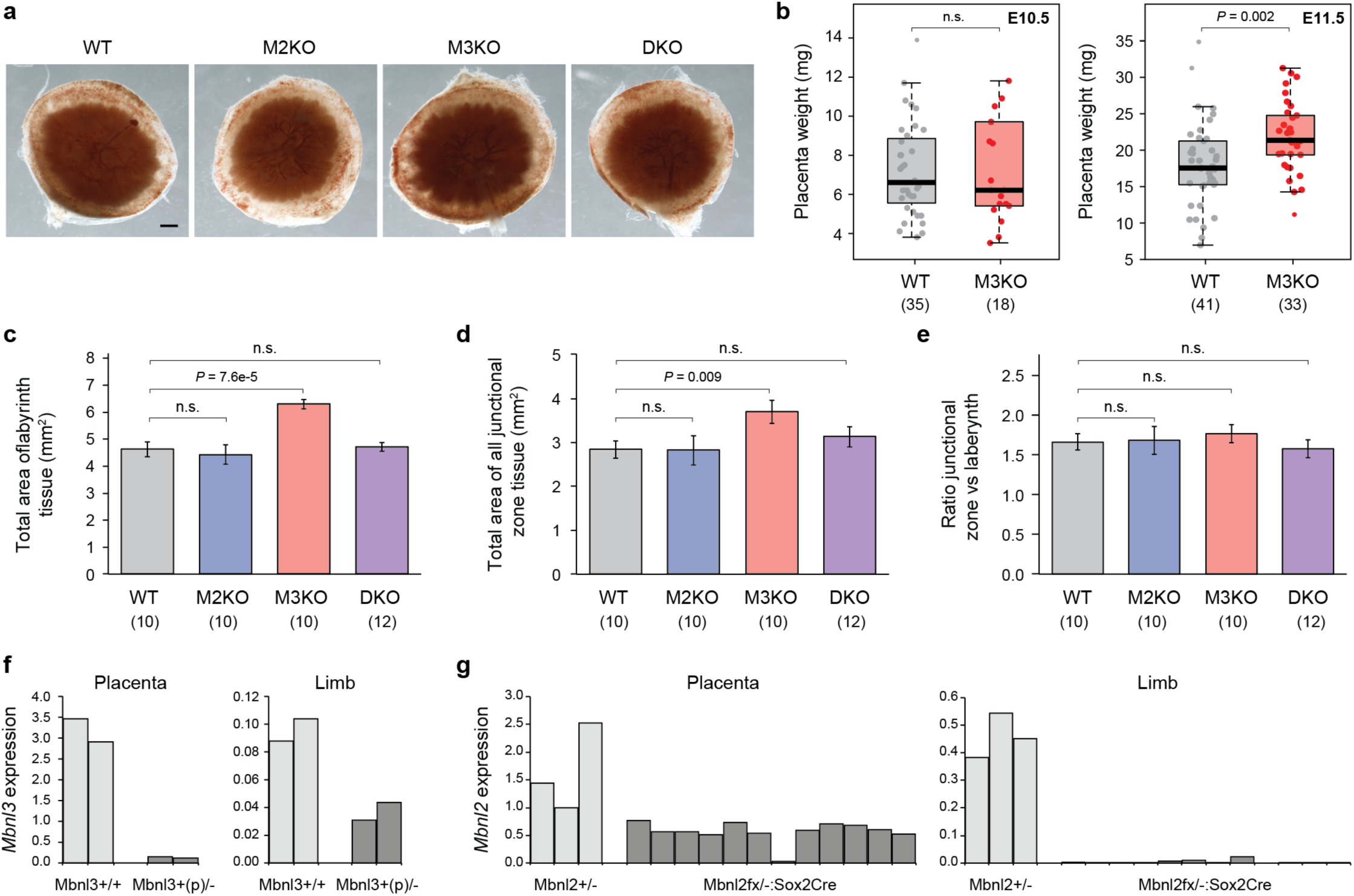
Mbnl3 restricts placental growth. **a**, Representative images of whole placentas at E18.5 for the indicated genotypes. **b**, Effect of *Mbnl3* knockout (M3KO) on placental weight at E10.5 and E11.5. **c-e**, Quantification of the total area of labyrinth tissue (c), total area of junctional zone tissue (d) and ratio between labyrinth and junctional zone tissues for the indicated genotypes. **f**,**g**, *Mbnl3* (f) and *Mbnl2* (g) expression quantified by qPCR in the indicated genotypes. Each bar corresponds to a single placenta/limb sample. For (b-e), the number of placentas/embryos analysed for each genotype are indicated in brackets and significance levels are calculated by Wilcoxon rank-sum tests.

**Extended Data Fig. 5.**
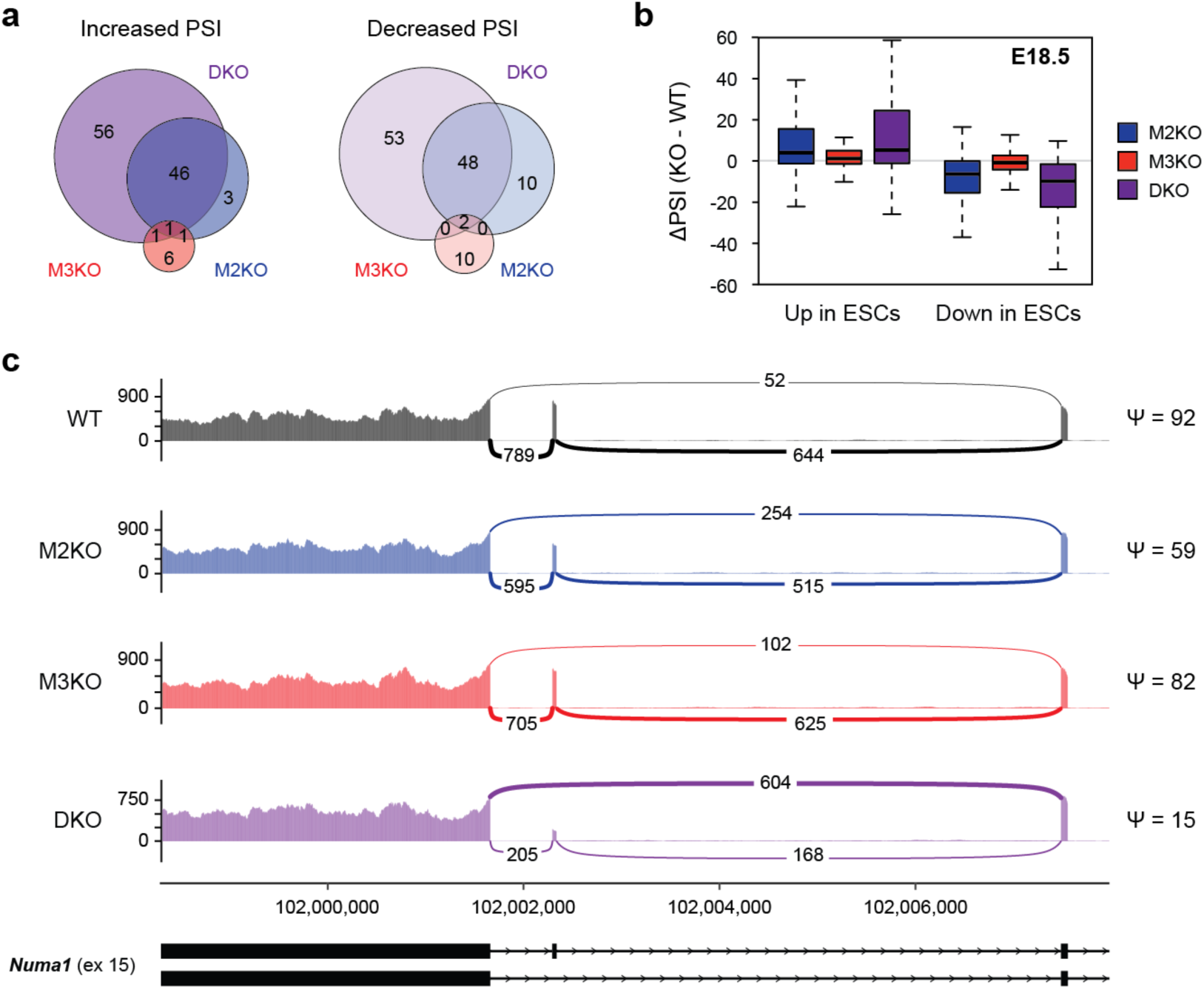
Effect of *Mbnl2, Mbnl3* and double *Mbnl2/3* knockout on alternative splicing. **a**, Venn diagrams showing the overlap in differential spliced exons between *Mbnl2* KO (M2KO), *Mbnl3* KO (M3KO) and double KO (DKO) placentas at E18.5. **b**, Boxplots showing the change in PSI in the indicated KO vs WT placentas at E18.5 for all exons with differential inclusion between embryonic stem cells and differentiated cell and tissues. **c**, Sashimi plots showing usage of *Numa* exon 15 in placentas with the indicated genotypes at E13.5. The PSI values for the exons in the different genotypes are shown to the right of the plots.

**Extended Data Fig. 6.**
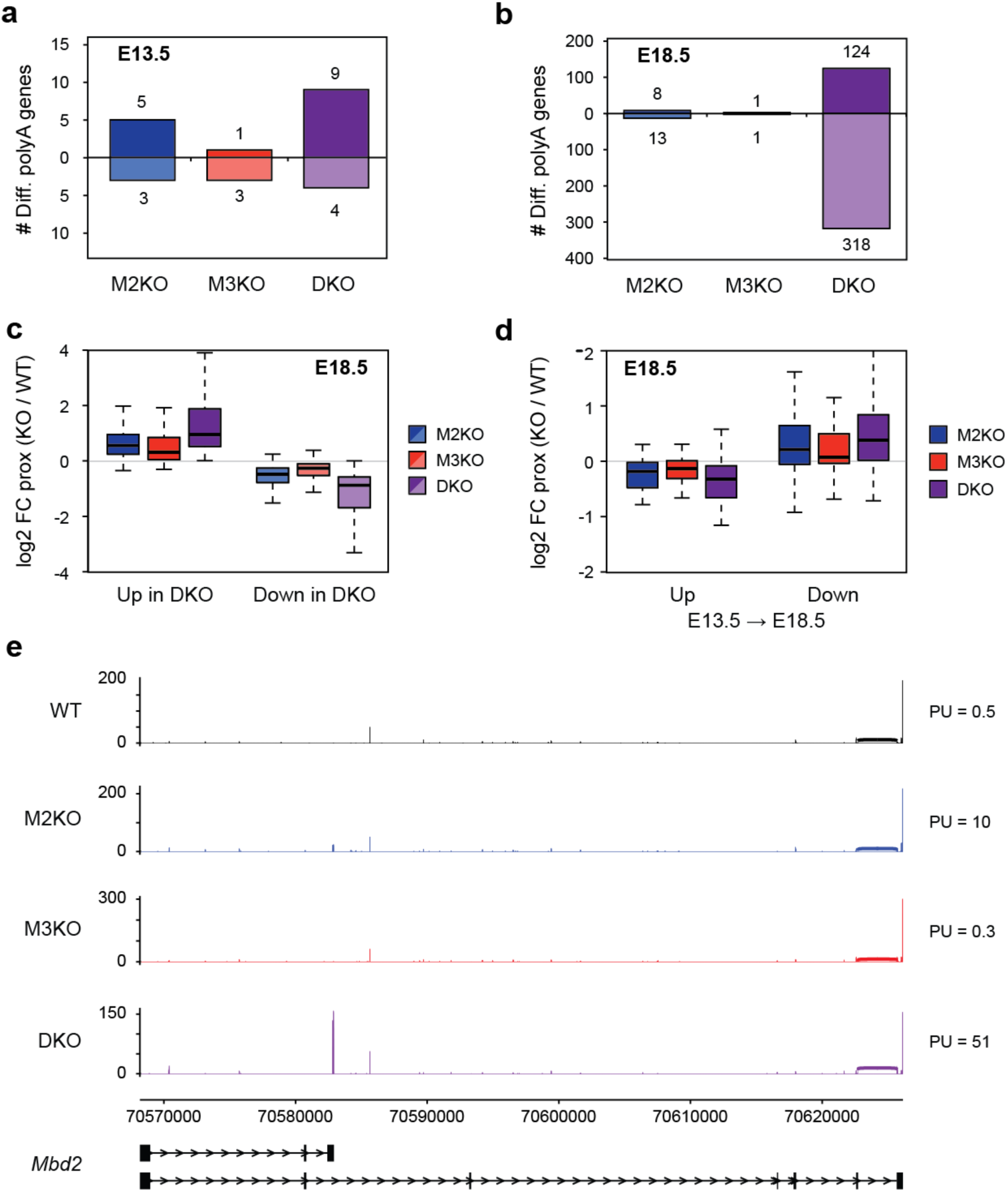
Effect of *Mbnl2, Mbnl3* and double *Mbnl2/3* knockout on alternative polyadenylation. **a**,**b**, Bar charts showing the number of genes with differentially used alternative polyadenylation (polyA) sites in *Mbnl2* KO (M2KO), *Mbnl3* KO (M3KO) and double KO (DKO) placentas at E13.5 (a) and E18.5 (b). Genes whose proximal polyA site increased usage in KOs are tallied above the x-axis whilst genes with reduced proximal polyA usage are tallied below. **c**, Boxplots showing the difference in proximal polyA site usage (log2 fold change [FC]) in the indicated KOs vs WT placentas for all genes with differentially used polyA sites in DKOs vs WTs at E18.5. Up in DKO, N = 124; down in DKO, N = 318. **d**, Boxplots showing the change in proximal polyA site usage in the indicated KO vs WT placentas at E18.5 for all genes polyA site pairs that are differentially used between E13.5 and E18.5 WT placenta. Up in E18.5, N = 31; down in E18.5, N = 77. **e**, Sashimi plots showing usage of the two competing polyA sites in *Mbd2* in placentas with the indicated genotypes at E13.5 based on 3’-seq data. The percent of usage (PU) values for the proximal polyA site in the different genotypes are shown to the right of the plots. Coordinates correspond to mm10.

**Extended Data Fig. 7.**
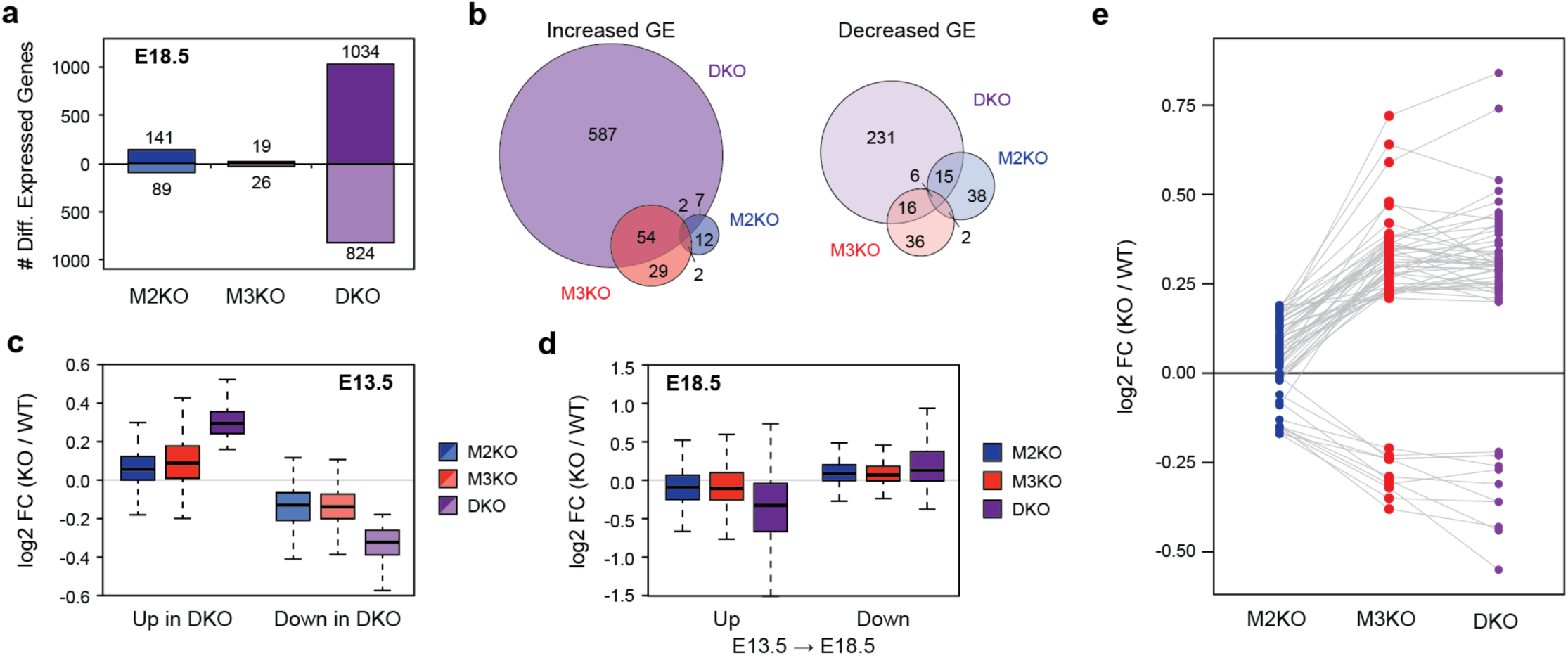
Effect of *Mbnl2, Mbnl3* and double *Mbnl2/3* knockout on gene expression. **a**, Bar charts showing the number of differentially expressed genes in *Mbnl2* KO (M2KO), *Mbnl3* KO (M3KO) and double KO (DKO) placentas at E18.5. Genes with increased expression in KOs are tallied above the x-axis whilst genes with reduced expression are tallied below. **b**, Venn diagrams showing the overlap in differentially expressed genes between M2KO, M3KO and DKO placentas at E18.5. **c**, Boxplots showing the difference in gene expression (log2 fold change [FC]) in the indicated KOs vs WT placentas for all genes differentially expressed in DKOs vs WTs at E13.5. **d**, Boxplots showing the change in expression in the indicated KO vs WT placentas at E18.5 for all genes differentially expressed between E13.5 and E18.5 WT placenta. **e**, Dot plots showing the change in expression in the indicated KO vs WT placentas at E13.5 for genes that are differentially expressed only in M3KOs and DKOs.

**Extended Data Fig. 8.**
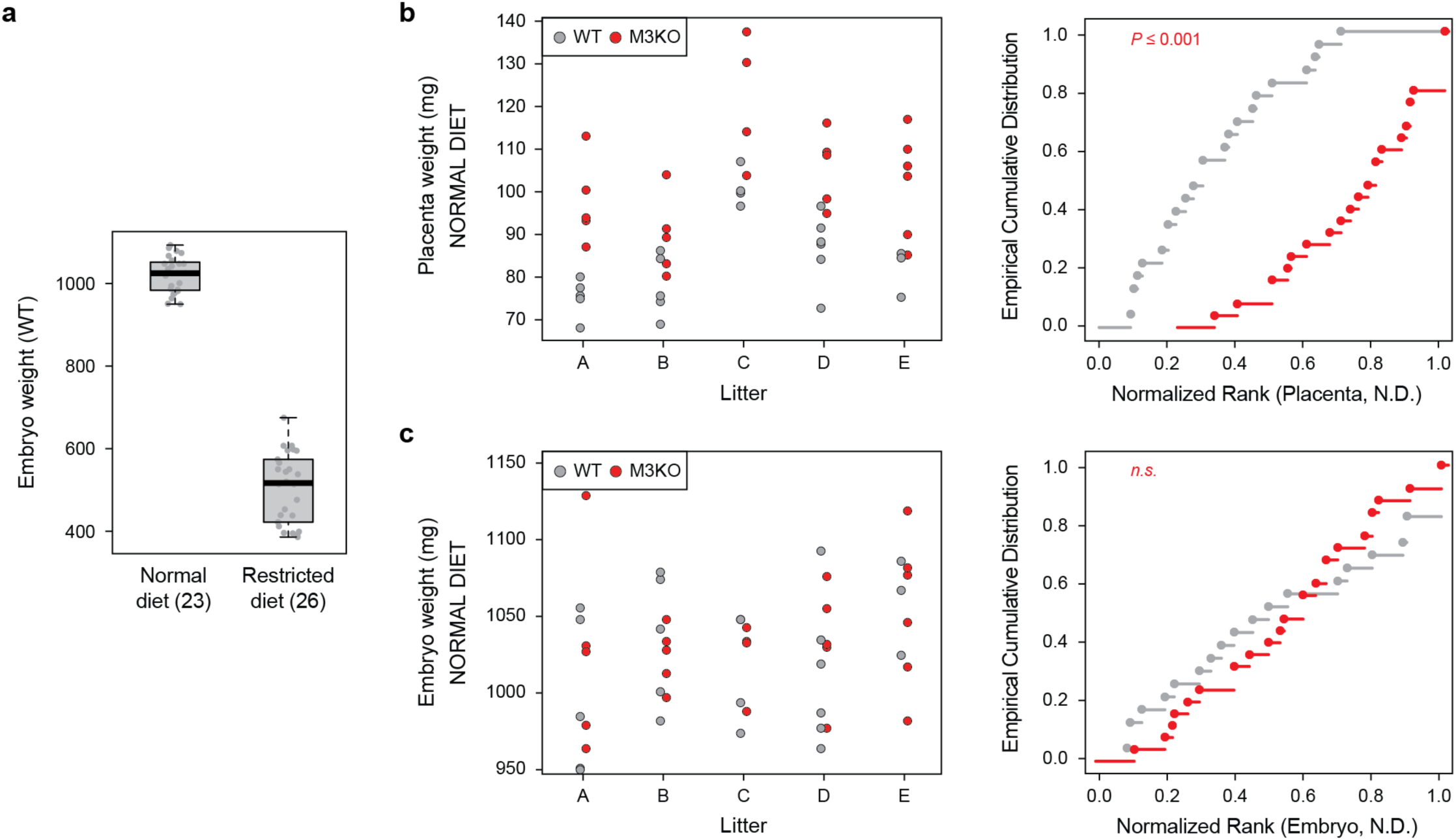
*Mbnl3* KO reduces the effect of mid-gestational maternal calorie restriction on fetal growth. **a**, Weight distributions of WT embryos under normal or calorie restricted diets. **b**,**c**, Dot plots showing the weights of individual WT (grey) and *Mbnl3* KO (red, M3KO) placentas (b) and embryos (c) harvested from five mothers fed with normal diet (A-E) and the corresponding empirical cumulative distribution plot for each genotype. Significance levels were calculated by a permutation test with 1000 iterations swapping the genotype labels within each litter.

