## Supplementary Materials for "The X-linked splicing regulator MBNL3 has been co-opted to restrict placental growth in eutherians"

### Supplementary Discussion

*Emergence of the short Mbnl3 isoform*

Intricately linked to placental recruitment of *Mbnl3* has been the emergence of a short isoform lacking the first zinc finger pair. Whilst alternatively spliced transcripts with the potential to code for a short isoform have been described for *Mbnl1* and *Mbnl2*, these appear to be poorly translated and to produce unstable proteins [1-3]. Indeed, production of these alternative transcripts has been proposed to act as a mechanism by which Mbnl genes can limit their own production, as the AS event generating them is auto-regulated by Mbnl proteins [1,4]. Therefore, the production of a stable and highly expressed short protein isoform lacking the first zinc finger pair is unique to eutherian *Mbnl3*. This further suggests that some of the amino acid changes seen in eutherian *Mbnl3* may be linked to stabilization of the truncated protein and/or enhanced short isoform translation.

The short isoform in placenta is mainly generated through the use of a novel eutherian-specific transcription start site. Interestingly, this transcription start site is closely associated with two eutherian-specific transposable elements of the MER53 and MER103c families, which are ~200 and ~850 bps away from the eutherian-specific TSS, respectively, and whose sequence appear largely conserved across this lineage. Given that transposable elements have been repeatedly shown to provide the raw evolutionary material for the evolution of new enhancers and transcription start sites [5,6], it is tempting to speculate their insertion in eutherian ancestors might be associated to the origin of the short Mbnl3 isoform in placenta.

*Evolution of the binding preferences of Mbnl3 isoforms*

The majority of full-length Mbnl proteins, including Mbnl1 and Mbnl2 from all tested species and Mbnl3 from non-eutherian species, have a strong GCUU binding preference (this paper and [7-10]). In contrast, we found the long isoform of mouse Mbnl3 alongside the majority of short Mbnl isoforms tested, including mouse Mbnl3, to preferentially bind a GCA/U motif and show weaker binding preferences. Previous studies have suggested the GCUU binding preference of canonical Mbnl proteins is largely driven by the first zinc finger pair and in particular ZF2 [11-13]. This is in keeping with our results for the short Mbnl isoforms, which lack this zinc finger pair, and suggests the binding preference changes seen for full-length eutherian Mbnl3 vs. other Mbnl proteins could be due to either a switch in dominance between the zinc fingers and/or changes in the binding preferences of ZF2. Furthermore, it raises the possibility that the that initial RNA interaction event(s) that mediated reduced placental growth were mediated by the second zinc finger pair and that there was then subsequent selection for mutations that favored these interactions. Our chimeric experiments revealed the binding preference of mouse Mbnl3 are determined by both the zinc finger domains, which are known to mediate binding, and the disordered linker region between the zinc finger domains. The precise function of the linker in altering the binding preference of Mbnl proteins is unclear; however, there is a growing pool of evidence that disordered regions can directly bind RNA and may cooperate with globular RNA motifs in this process [14]. Indeed, previous studies have found the linker to play an important role in RNA binding [15-17]. Alternatively, the linker could directly or indirectly regulate the accessibility/availability of individual zinc-fingers for binding.

**Supplementary Tables**

**Supplementary Table 1 - RNA-seq data used in this study.** For each species, SRA identifiers, read number, read length and source are provided for each public RNA-seq file used in this study. Groups used to identify placenta-enriched factors are also provided. For RNA-seq samples generated in this study, mapping statistics are also provided.

**Supplementary Table 2 - Mouse splicing factors enriched in trophoblastic tissues.** log2 fold changes and expression (log10TPM) in the trophoblastic/placenta tissue is provided for mouse splicing factors for each comparison: (i) placenta vs tissues, (ii) trophectoderm (TE) vs inner cell mass (ICM), and (iii) trophoblast stem cells (TS) vs embryonic stem cells (ESC) and extraembryonic endoderm stem cells (XEN). Only genes with a minimal expression of TPM $\geq$ 10 in the target tissue are included.

**Supplementary Table 3 - Human splicing factors enriched in placenta.** log2 fold change between placenta and non-placental tissues and expression (log10TPM) in the placenta tissue is provided for human splicing factors with an expression in placenta of TPM $\geq$ 10.

**Supplementary Table 4 - Cow splicing factors enriched in placenta.** log2 fold change between placenta and non-placental tissues and expression (log10TPM) in the placenta tissue is provided for cow splicing factors with an expression in placenta of TPM $\geq$ 10.

**Supplementary Table 5 - Opossum splicing factors enriched in placenta.** log2 fold change between placenta and non-placental tissues and expression (log10TPM) in the placenta tissue is provided for opossum splicing factors with an expression in placenta of TPM $\geq$ 10.

**Supplementary Table 6 - Usage of transcription start sites in eutherian *Mbnl3*.** For each sample and species, number of reads and percent over the total mapping to each competing splice junction are provided. Competing junctions consist of alternative splice donors (normally resulting from alternative promoters) joint to the same splice acceptor in the third ancestral exon (see scheme in Fig. 2a). Each donor annotated in *vast-tools* is numbered, starting from 0 and the coordinate on the X-chromosome is provided in the header (e.g. 3:132413583; donor 3, coordinate chrX:132413583). The coordinates and donor numbers are depicted in Extended Data Fig. 2a-c for each species. The color code for each column corresponds to the scheme of Fig 2a: Black, canonical long isoform using the first ATG-encoding exon; Grey, ancestral transcription start site (TSS) skipping the first ATG-encoding exon; Red, eutherian specific TSS; orange, rodent or primate specific TSS. The column "Group" shows the sample groupings using to generate the quantifications displayed in Fig. 2c and Extended Data Fig. 2e,f.

**Supplementary Table 7 - Differentially spliced exons.** Exons found to be differentially spliced in at least one KO vs WT comparison at the E13.5 or E18.5 stage. Gene ID, gene symbol, VastID, genomic coordinate (mm10) and exon length are provided alongside the inclusion level (PSI) for each WT and KO replicate and the change in developmental time (PSI_E18.5 - PSI_E13.5). Key: the exon is upregulated (U), downregulated (D) or not differentially spliced (N) in the KO vs WT comparison for the double KO, *Mbnl2* KO and *Mbnl3* KO (e.g. DDN, downregulated in double KO and *Mbnl2* KO but not *Mbnl3* KO).

**Supplementary Table 8 - Enriched Gene Ontology terms for genes with differentially spliced exons.** Significantly enriched terms for each KO vs WT comparison at E13.5 or E18.5 as provided by DAVID. Only GOTERM DIRECT (biological process, molecular function and cellular component) and KEGG pathways were tested.

**Supplementary Table 9 - Pairs of alternative polyA site analyzes in this study.** For E13.5 or E18.5, a pair of alternative polyA sites per gene was selected (see Methods for details). Genomic features, raw read counts, and log2 fold changes for each KO vs WT (KO/WT) comparison and E18.5 vs E13.5 using the proximal polyA site as reference are provided. Enhanced/Repressed indicates a given polyA pair is significantly differentially regulated in the KO and the direction refers to the proximal polyA. E.g. an enhanced pair means the proximal site is upregulated in the KO. KO_Up/KO_Down indicates that the proximal site is up/down-regulated in the KO, but not reaching statistical significance.

**Supplementary Table 10 - Differentially expressed genes.** For each stage and KO comparison, the log2 fold change and FDR-adjusted p-value (padj) are provided for each expressed gene. Mbnl3-specific genes at E13.5 are defined as those differentially expressed in the *Mbnl3* KO and double KO comparisons but not in the *Mbnl2* KO one. Genes with a potential role in nutrient uptake are highlighted in green.

**Supplementary Table 11 - Enriched Gene Ontology terms for differentially expressed genes.** Significantly enriched terms for up- and down-regulated genes for each KO vs WT comparison at E13.5 or E18.5 as provided by DAVID. Only GOTERM DIRECT (biological process, molecular function and cellular component) and KEGG pathways were tested.

**Supplementary Table 12 - Enriched Gene Ontology terms and EnrichR results for differentially expressed genes at E11.5.** Significantly enriched terms for up- and down-regulated genes in *Mbnl3* KO vs WT at E11.5 as provided by DAVID are shown. Only GOTERM DIRECT (biological process, molecular function and cellular component) and KEGG pathways were tested. The results for the regulatory enrichment using EnrichR are also provided for upregulated genes.

**Supplementary Table 13 - Splicing regulators.** List of 197 splicing regulators used in this study and Ensembl IDs for one-to-one orthologs in human, mouse, cow and opossum.

**Supplementary Table 14 - Primers used in this study.** Restriction sites added to primers and used in cloning are highlighted in red (enhancer fragments #1, SacII and NotI, #2, SalI and SalI, #3 NotI and NotI. Mbnls for RNA compete, AscI and SbfI).

**Supplementary Table 15 - Config file for calling ESC-differential exons.** Config file used to run Get_Tissue_Specific_AS.pl to calculate ΔPSI values (ESC - other cell/tissue types).
